# Genomic environments scale the activities of diverse core promoters

**DOI:** 10.1101/2021.03.08.434469

**Authors:** Clarice KY Hong, Barak A Cohen

## Abstract

A classical model of gene regulation is that enhancers provide specificity while core promoters provide a modular site for the assembly of the basal transcriptional machinery. However, examples of core promoter specificity have led to an alternate hypothesis in which specificity is achieved by core promoters with different sequence motifs that respond differently to genomic environments containing different enhancers and chromatin landscapes. To distinguish between these models, we measured the activities of hundreds of diverse core promoters in four different genomic locations and, in a complementary experiment, six different core promoters at thousands of locations across the genome. While genomic locations had large effects on expression, the intrinsic activities of different classes of promoters were preserved across genomic locations, suggesting that core promoters are modular regulatory elements whose activities are independently scaled up or down by different genomic locations. This scaling of promoter activities is non-linear and depends on the genomic location and the strength of the core promoter. Our results support the classical model of regulation in which diverse core promoter motifs set the intrinsic strengths of core promoters, which are then amplified or dampened by the activities of their genomic environments.

## Introduction

In the classical model of gene regulation, the core promoter serves as a universal platform for the assembly of the basal transcriptional machinery, while the specificity of expression is provided by distal enhancers and the chromatin landscape. However, some examples of core promoter specificity seem to challenge this model. Several studies suggest that different core promoters are specific for distinct sets of enhancers (Gehrig et al. 2009; Li and Noll 1994; Merli et al. 1996; Sharpe et al. 1998), and can even trap different enhancers at the same genomic location (Butler and Kadonaga 2001). Some transcription factors also preferentially activate core promoters containing specific motifs (Emami et al. 1995; Juven-Gershon et al. 2006; Parry et al. 2010; Haberle et al. 2019). More recently, a genome-wide massively parallel reporter assay (MPRA) showed that housekeeping and developmental core promoters respond to distinct classes of enhancers (Zabidi et al. 2015; Arnold et al. 2017), arguing that enhancer-promoter compatibility contributes to specificity in the genome. These data have led to an alternate model, which we refer to as the ‘promoter compatibility’ hypothesis, in which core promoters with different sequence elements respond specifically to the enhancers and chromatin features that comprise distinct genomic environments. Determining whether the specificity of gene expression is governed by enhancers and chromatin features or by enhancer-promoter compatibility is crucial to understanding a variety of biological processes including cell-type specific regulatory programs and models of gene evolution.

The core promoter is the ∼100bp region around the transcription start site and is responsible for accurately position RNA polymerase II and binding general transcription factors (Roy and Singer 2015; Haberle and Stark 2018). It is now known that core promoters are a diverse set of sequences containing specific DNA sequence motifs, also known as core promoter elements or motifs. The most well-known core promoter motif is the TATA box, however, the TATA box is only present in 10-20% of metazoan core promoters (Yang et al. 2007), suggesting that other motifs might have evolved for different functions. The different motifs have been associated with different functions, for example, the TATA box is often enriched in developmental promoters and show a ‘sharp’ pattern of transcription initiation, while high CpG promoters tend to contain other less well-characterized motifs and is thought to be associated with a broader pattern of transcription initiation (Lenhard et al. 2012).

A strong prediction of the promoter compatibility hypothesis is that the relative strengths of different core promoters will change at different genomic locations because the distal enhancers and chromatin environments at different locations will be compatible with different types of core promoters (Fig. 1A). Here, we tested the promoter compatibility hypothesis by assaying hundreds of diverse core promoters at four different genomic locations and further extend our results genome-wide by assaying six core promoters across thousands of genomic locations. We find that the intrinsic activities of core promoters are preserved across genomic locations, suggesting that diverse core promoter motifs do not interact specifically with genomic environments, a finding consistent with the classical model of gene regulation. Different core promoter motifs set the intrinsic strength of the promoter, and those intrinsic strengths are independently scaled up or down by the genomic environment.

**Fig. 1.**
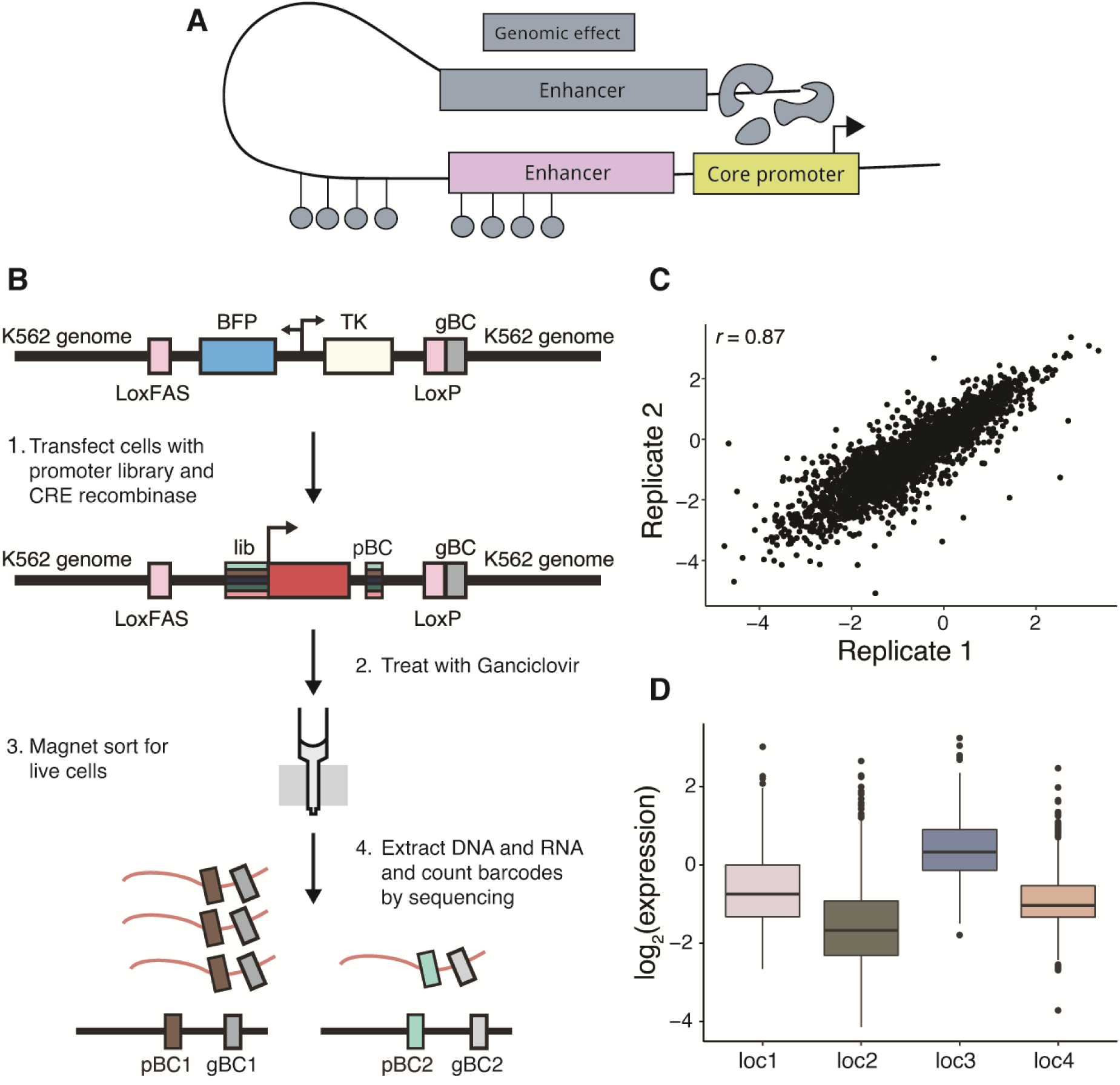
Measurements of core promoter library at four genomic locations by patchMPRA. **(A)** Schematic of gene regulation by the core promoter, adjacent *cis*-regulatory sequences and the genomic environment. **(B)** Schematic of patchMPRA method (see Methods for details). BFP: blue fluorescent protein; TK: thymidine kinase; gBC: genomic barcode; pBC: promoter barcode. **(C)** Reproducibility of core promoter measurements from independent patchMPRA transfections. **(D)** The expression of all core promoters in the library at each genomic location.

## Results

### Measurement of diverse core promoter activities at different genomic locations

We first created a library of reporter genes driven by diverse core promoters. The library contains 676 133bp core promoters spanning a variety of promoter features from Haberle *et al*. (Haberle et al. 2019), including the most common mammalian core promoter motifs (TATA, DPE and TCT), CpG islands and housekeeping (hk) and developmental (dev) promoters that do not contain any known core promoter motifs (Supplemental Table S1 and S2). To provide redundancy in the measurements, we included ten copies of each individual core promoter in the library, each with a unique barcode (promoter BC; pBC) in the 3’ UTR. Because basal expression of the core promoters was expected to be weak, we included a common proximal enhancer directly upstream of the core promoters to boost expression (Methods).

Using patchMPRA (<u>pa</u>rallel targeting of <u>ch</u>romosome positions by <u>MPRA</u>), we measured the expression of the core promoter library in parallel at four genomic locations previously shown to have diverse expression levels and chromatin marks in K562 cell lines (Supplemental Table S3 and Supplemental Fig. S1) (Maricque et al. 2019). Each cell line contains a single ‘landing pad’ at a different genomic location. Each landing pad has a unique genomic barcode (gBC) indicating its location in the genome and a pair of asymmetric Lox sites to facilitate site-specific recombination of the library. We pooled the four landing pad lines and integrated the library into the cells by cotransfection with CRE recombinase (Maricque et al. 2019). When a library member recombines into a landing pad it produces a transcript with two unique barcodes in its 3’ UTR; a pBC specifying the core promoter and a gBC indicating its genomic location. By tabulating the pBC-gBC pairs in the mRNAs from the pool we obtained expression measurements for every core promoter at each genomic location in parallel (Fig. 1B).

We obtained reliable measurements of every core promoter at all four genomic locations. We recovered 70-80% of all promoter barcodes and 99% of all promoters at all landing pads (Supplemental Figs. S2A and S2B). The three biological replicates showed high reproducibility (average Pearson’s *r* = 0.87) (Fig. 1C and Supplemental Fig. S2C) and the locations of the landing pad had large effects on library expression that were consistent with previous studies (compare Fig. 1D to Supplemental Fig. S3A; (Maricque et al. 2019) indicating that the genomic environment is not drastically altered by a diverse core promoter library. The data allowed us to compare the effects of the four genomic environments on the different classes of core promoters.

### The effects of genomic locations on core promoters

The promoter compatibility hypothesis predicts that the same genomic environment will impact different classes of promoters differently. In contrast to this prediction, the genomic effect was similar on all promoter classes: more permissive genomic locations boosted the expression of all promoter classes regardless of their motif composition or their hk or dev designation (Fig. 2A). However, the magnitude of the genomic effect is not the same for all promoter classes. To quantify the contribution of the genomic location and core promoters to gene expression we performed ANOVA on each class of promoters. In general, genomic locations have a larger effects on dev promoters than hk promoters regardless of their motif composition (Fig. 2B). Thus, we did not distinguish between the motif classes and focused on the hk and dev groupings for downstream analysis.

**Fig. 2.**
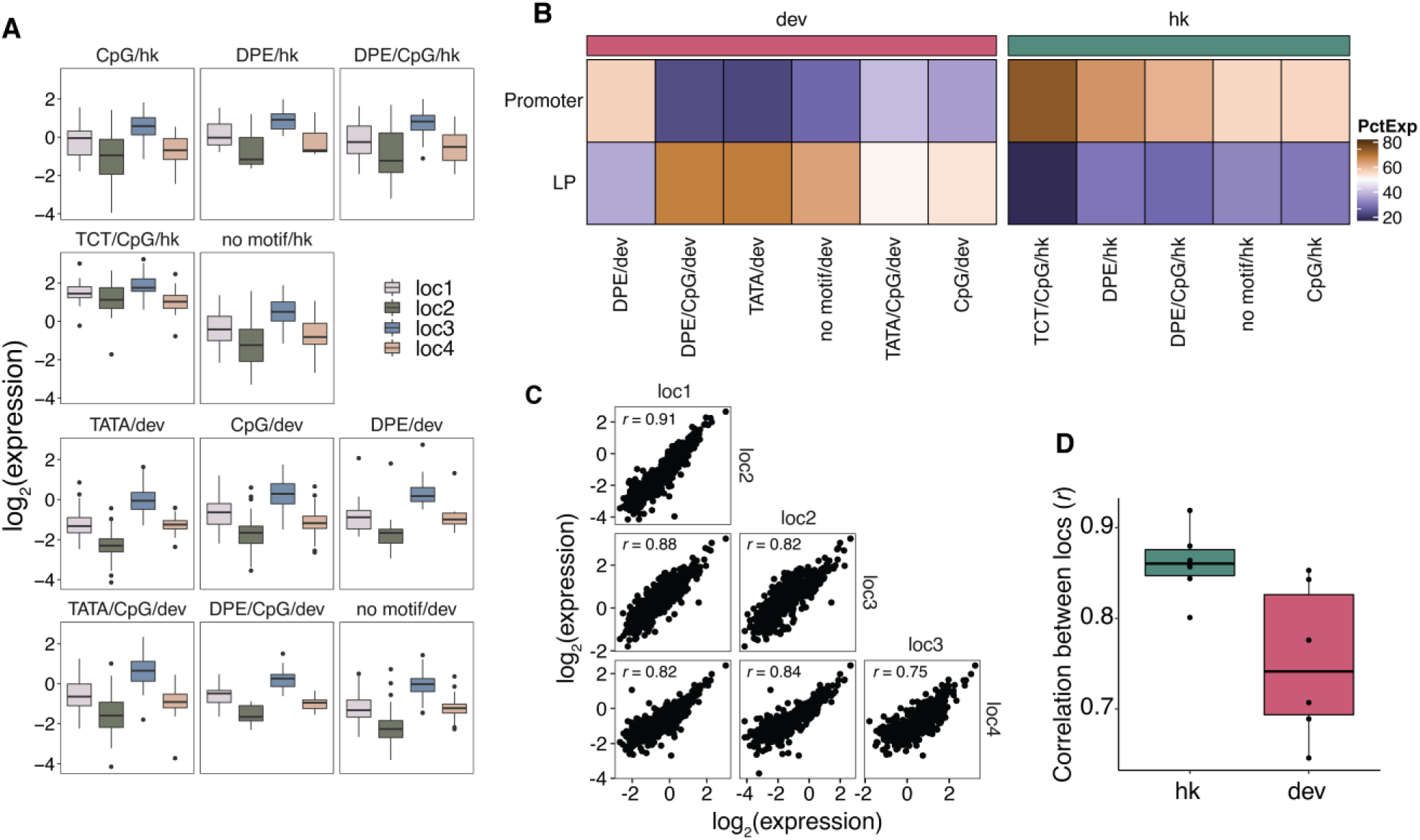
Effects of genomic locations on core promoter activity. **(A)** Expression of each class of core promoter motifs at each genomic location. **(B)** Amount of variance explained by core promoter and genomic location respectively using linear models fit on each class of core promoters separately. **(C)** Pairwise correlations (Pearson’s *r*) of core promoter activity between the different genomic locations. **(D)** All pairwise correlations (Pearson’s *r*) between genomic locations for hk and dev core promoters.

We further examined whether hk and dev core promoter activities are scaled by different genomic environments. We define scaling as the degree to which core promoter activities correlate between genomic locations. High correlations between genomic locations indicate that the rank order of core promoter activities is preserved across genomic locations. While promoter activities were highly correlated between genomic locations regardless of the class of promoter (Pearson’s *r* = 0.74 - 0.9, Spearman’s ρ = 0.72 - 0.88) (Fig. 2C), dev promoters were consistently less correlated than hk promoters (Fig. 2D). Dividing the promoters into classes containing different motifs showed that each class also had substantial differences in correlations between genomic locations (Supplemental Fig. S3B). These results do not depend on the proximal enhancer immediately adjacent to the core promoter used to boost expression because replicate experiments at locations 1-3 without the enhancer yielded similar results (Supplemental Figs. S4A-C). The expression of libraries with and without the proximal enhancer is also largely correlated at locations 1-3 (Supplemental Fig. S4D), which suggests that scaling by different genomic locations does not depend on the proximal enhancer. Taken together these results suggest that genomic environments scale the activities of all core promoters, but that the quantitative extent of scaling can differ between promoter classes.

### Intrinsic promoter strength explains differences between promoter classes

A striking difference between hk and dev promoters in our library is that they have different mean levels of expression—hk promoters are consistently stronger than dev promoters at all genomic locations (Fig. 2A and Supplemental Fig. S5A). Thus, any differences between hk and dev promoters might be confounded by their difference in strength. To test if strength explains the differences between hk and dev promoters, we divided all core promoters into strong or weak bins based on their strengths and sampled equal numbers of hk and dev promoters within each bin to avoid confounding the results by hk/dev class. Plotting the effect of genomic position on strong and weak promoters showed that the direction of the effect was the same, but that there were larger differences between genomic locations for weak promoters (Fig. 3A). We quantified the contributions of genomic locations and promoters within strong and weak bins respectively and found that the genomic environment has a larger impact on weak promoters compared to strong promoters (Fig. 3B). For strong promoters, genomic environments and core promoters contribute almost equally to gene expression (∼42% and ∼46% respectively), but for weak promoters, genomic environments contribute ∼73%, while core promoters contribute only ∼15%. Weak promoters are also consistently less correlated than strong promoters (Fig. 3C). Again, assaying the library without an upstream proximal enhancer at locations 1-3 showed similar results (Supplemental Figs. SB5 and S5C). Finally, we sampled a set of hk and dev promoters with similar average strengths (Supplemental Fig. S5D) and compared their correlations across genomic locations. Using only this subset of promoters, correlations across genomic locations are comparable between hk and dev promoters (Supplemental Fig. S5E). The differences in how genomic locations scale the activities of each core promoter subclass is also largely explained by the average strength of each promoter class (Supplemental Fig. S5F). These data show that the observed differences between different promoter classes is a consequence of promoter strength rather than a feature of the hk/dev distinction, indicating that the strength of a promoter is a key determinant of its interactions with the genomic environment.

**Fig. 3.**
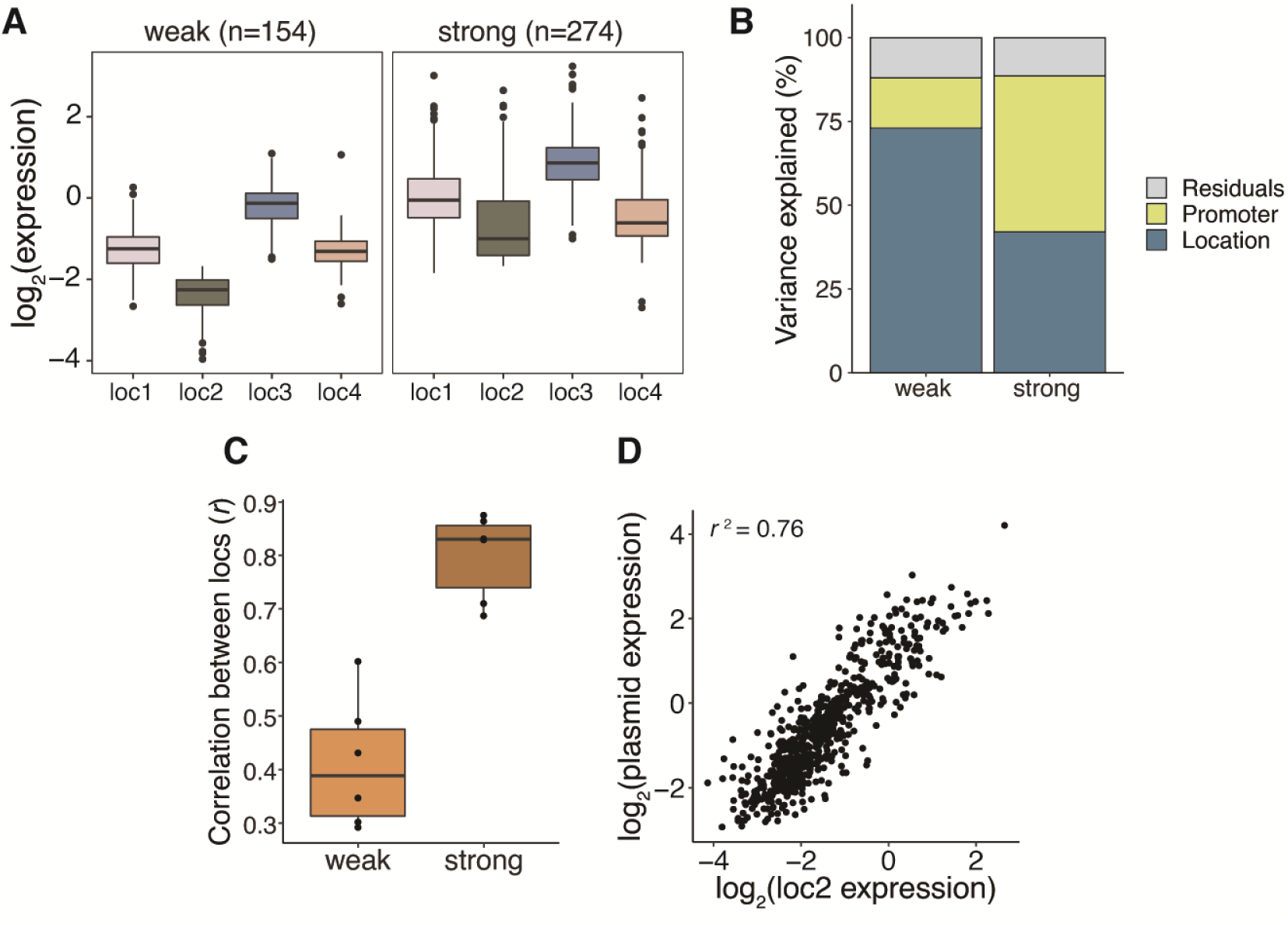
Intrinsic promoter strength explains differences between promoter classes. **(A)** The effect of genomic location on the expression of weak and strong core promoters. **(B)** Amount of variance explained by core promoters and genomic locations respectively using linear models fit on weak and strong promoters separately. **(C)** All pairwise correlations (Pearson’s *r*) between genomic locations for weak and strong core promoters. **(D)** Correlation (Pearson’s *r*) between promoter activity measured on plasmids and promoter activity at loc2.

Given the importance of the interaction between promoter strength and genomic location, we next asked if core promoter strengths, as measured in the genome, reflect the promoters’ intrinsic activities. If this is true, then the measurements in the genome should correlate with measurements on plasmids, assuming that plasmids represent a neutral environment that reflect the intrinsic activities of core promoters. Thus, we performed an episomal MPRA on the core promoter library in K562 cells. The plasmid measurements are well-correlated with expression at each genomic location (Pearson’s *r*^2^ = 0.59-0.76; Fig. 3D and Supplemental Fig. S6A), indicating that the relative intrinsic activities of core promoters are preserved when integrated into the genome. We were also able to predict activity in the genome using activity on plasmids (adjusted *r*^2^ = 0.78; Supplemental Fig. S6B). These results demonstrate that genomic locations scale the intrinsic activities of strong and weak promoters to different extents, suggesting that the main role of diverse core promoter motifs is to set the intrinsic strength of the promoter rather than direct specific interactions with the genomic environment.

### Core promoter scaling is a genome-wide phenomenon

To extend the results we observed at four genomic locations to diverse locations across the genome, we selected six core promoters (three hk and three dev) spanning a range of expression levels and motifs within each class (Supplemental Table S4). We then measured their activities genome-wide using the TRIP (<u>T</u>housands of <u>R</u>eporters <u>I</u>ntegrated in <u>P</u>arallel) assay (Akhtar et al. 2013) in K562 cells (Supplemental Fig. S7A). Each core promoter was fused to a unique barcode (pBC) in its 3’UTR and cloned upstream of a reporter gene into a PiggyBac transposon vector for random delivery into the genome. No upstream proximal enhancer was included in these constructs. TRIP libraries were generated by incorporating >10^5^ random barcodes (tBCs) onto each core promoter reporter plasmid. After transposition, every genomic integration contains a unique pBC and tBC pair specifying the identity of the core promoter and its location in the genome respectively. This double barcoding strategy allowed us to pool promoter libraries into a single TRIP experiment. The replicates were highly correlated (Pearson’s *r*^2^ = 0.96, Supplemental Fig. S7B). In total, we mapped 41,083 unique integrations in the genome, ranging between 6078-7418 integrations per promoter (Supplemental Table S4 and S5).

Genomic positions have large effects on core promoter activities, with expression ranging more than 1000-fold for the same promoter across genomic locations (Fig. 4A). However, even with these large effects of genomic location, the rank order of promoter strengths is preserved across locations and correlates with mean expression in the landing pads (Fig. 4B and Supplemental Fig. S7C), which suggests that the effect of different genomic locations is to scale intrinsic promoter activities. Because PiggyBac is known to have a preference for H3K27-acetylated regions (Yoshida et al. 2017; Moudgil et al. 2020), we grouped the integrations by their locations into three groups: H3K27ac regions (n=14,275), within 50kb of a H3K27ac region (n=21,623) or far away from H3K27ac regions (n=5185) (Supplemental Fig. S7D). Integrations that are far from H3K27ac regions are generally weak, consistent with the idea that these locations are less permissive for expression. However, the rank order of promoters in these regions is the same as integrations in the other locations. Furthermore, integrations in or near H3K27ac regions span the entire >1000-fold dynamic range of our library, suggesting that there is still substantial diversity within H3K27ac regions. Taken together, these data indicate that core promoters are scaled by diverse genomic environments.

**Fig. 4.**
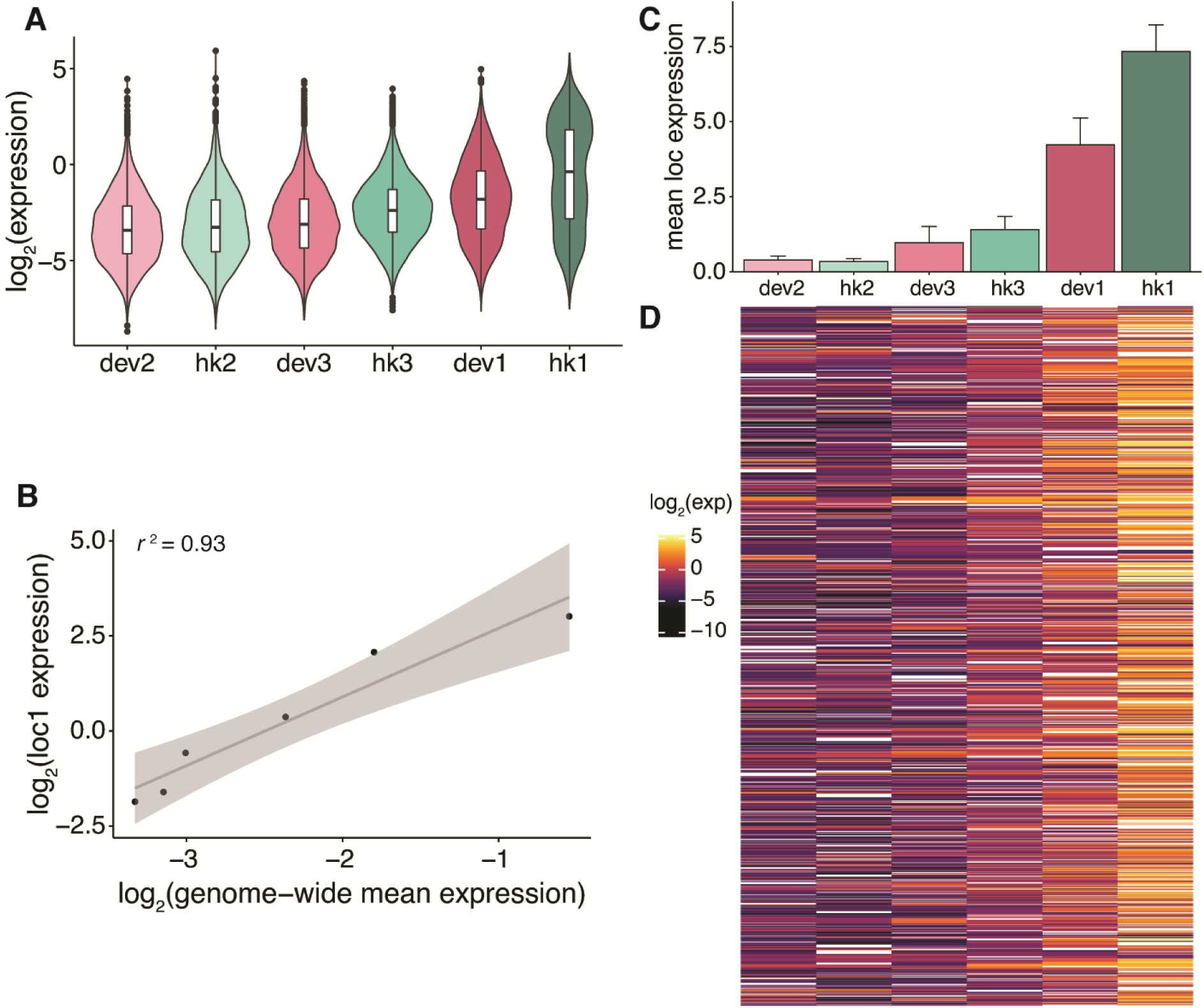
Core promoter scaling is a genome-wide phenomenon. **(A)** Expression of each core promoter across all mapped genomic locations sorted by increasing means measured by TRIP. Blue-green denotes hk promoters and pink denotes dev promoters. **(B)** Correlation (Pearson’s *r*) between mean expression of each core promoter genome-wide (measured by TRIP) and loc1. The shaded region around the fitted line represents the 95% confidence interval. **(C)** Mean expression of each core promoter from four genomic locations as measured by patchMPRA. Error bars represent the SEM. **(D)** Heatmap of expression of each core promoter (column) at each genomic region (row) that has ≥4 different integrated promoters. White boxes represent NA values.

To compare different promoters in the same genomic environment, we identified 1278 genomic regions in which at least 4 of the 6 promoters had integrated <5kb from each other (in separate cells) (Supplemental Table S6). These genomic regions are located across the entire genome and span diverse chromHMM annotations (Ernst and Kellis 2010; Ernst et al. 2011) (Supplemental Figs. S8A and S8B). Across these locations, expression consistently increases from the weakest (dev2) to strongest (hk1) promoter (Figs. 4C and 4D), showing that the relative strengths of core promoters are preserved across >1000 genomic locations with 1000-fold differences in expression. The expression of the promoters in each region also correlates well with expression in the landing pads, with >60% of locations having *r* > 0.7 (Supplemental Fig. S8C), and a linear model assuming independent effects of genomic region and promoter explains ∼54% of the variance in the data (Supplemental Fig. S8D). Thus, measurements of integrated promoters across diverse genomic positions demonstrates that core promoter scaling is a genome-wide phenomenon.

### Non-linear scaling of core promoters by genomic environments

We next explored the relationship between core promoter strength and genomic environments in the TRIP data. We ranked the TRIP genomic regions based on mean promoter expression and plotted the expression of the promoters (Fig. 5A). As expected, all six core promoters increase expression as genomic environments become more permissive. However, the rates at which their expression changes are different for strong and weak promoters. In less permissive regions, strong promoters increase rapidly, but then level off in more permissive regions. In contrast, weak promoters increase slowly in less permissive regions and then sharply in more permissive regions. To ensure that hk1 expression in activating regions is not saturated due to the dynamic range limits of TRIP, we tested hk1 with an upstream enhancer and it was expressed at still higher levels (Supplemental Fig. S8E). Thus, promoters with different strengths do not respond to differences in genomic environments in the same way.

**Fig. 5.**
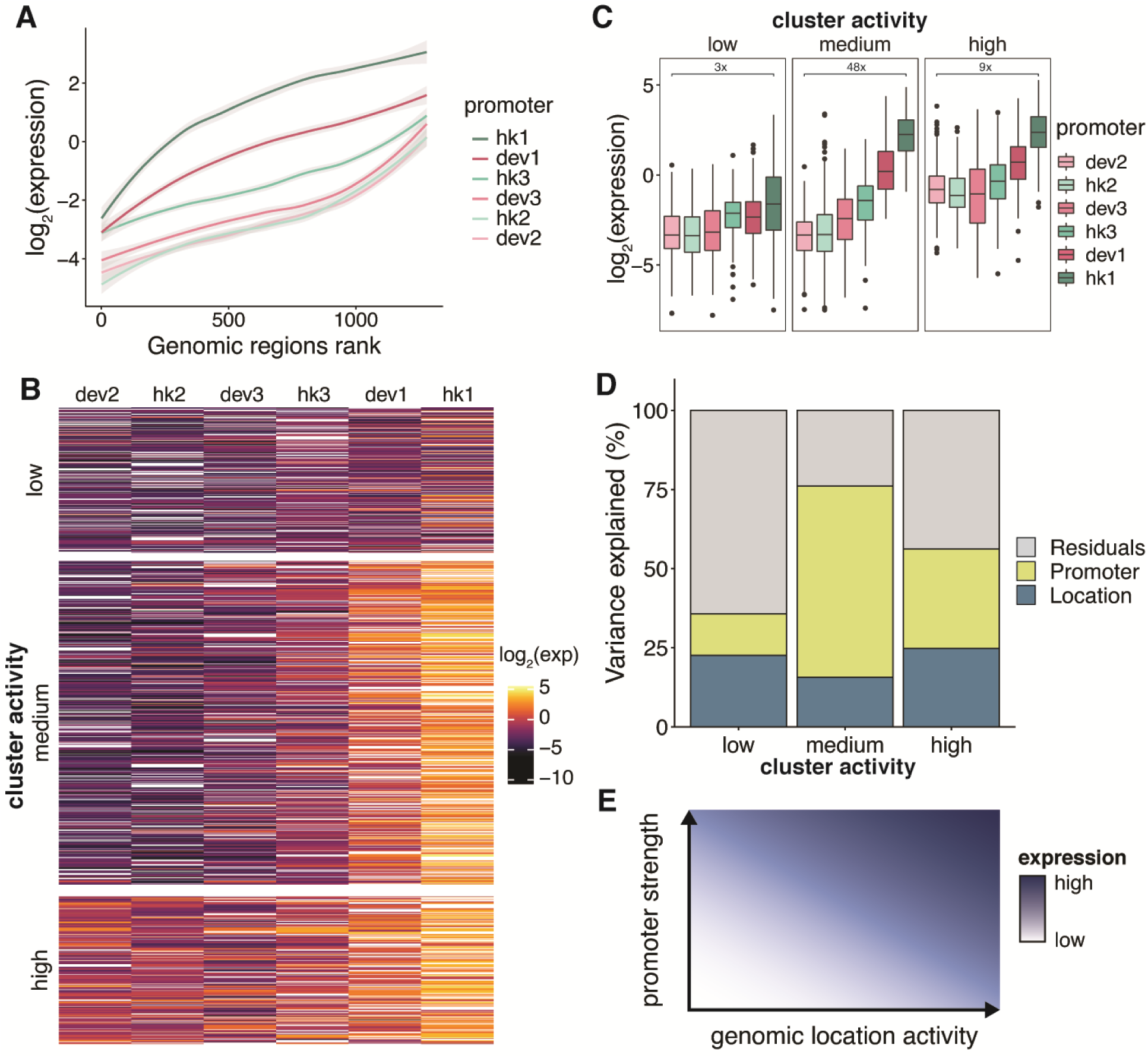
Non-linear scaling of core promoters by genomic environments. **(A)** Genomic regions defined by TRIP were sorted by the mean expression of the promoters in each region. The shaded region around the fitted line represents the 95% confidence interval. **(B)** Heatmap in Fig. 3b split into 3 clusters by k-means clustering. Clusters were assigned different activity levels based on the overall expression in the cluster. **(C)** Expression of core promoters in each genomic cluster. **(D)** Amount of variance explained by core promoters and genomic locations respectively using linear models fit on each genomic cluster respectively. **(E)** Summary model of the relationship between core promoter strength and genomic environment activity.

Interestingly, the curves in Fig. 5A separate by the intrinsic strength of the core promoters and not by their hk or dev identity. To emphasize this point we calculated the correlations between the curves of each promoter and show that the promoters cluster based on their intrinsic strengths, with the stronger promoters (dev1 and hk1) in one cluster and the others in another (Supplemental Fig. S9A). Integrations within 5kb of endogenous hk or dev promoters in K562 also showed no preference for hk or dev promoters respectively (Supplemental Fig. S9B). This result again highlights that a promoter’s strength, not class, determines its interaction with genomic environments.

The differences in the way core promoters respond to genomic environments in Fig. 5A also demonstrate that genomic environments do not scale promoter activities linearly. Although the rank order of core promoters is preserved across the genome, the fold change between strong and weak core promoters is different in different parts of the genome. To quantify the effects of different genomic environments, we identified three clusters of TRIP genomic regions that appear to have different levels of activity (Fig. 5B). While the clusters are defined by their average differences in core promoter expression, the extent of scaling is also different in each cluster (Fig. 5C). This difference in scaling is due to differences in the contributions of genomic location and promoter effects in the three clusters. In regions of the genome with low activity, genomic location contributes ∼23% to gene expression while core promoters contribute only ∼12%. In the cluster with high activity, genomic location also contributes about ∼24%, but core promoters contribute ∼31%, suggesting that differences in expression at these locations depend more on core promoter strength. In the cluster with medium activity, the core promoter contribution is much larger, explaining ∼64% of the variance compared to ∼16% by genomic location (Fig. 5D). Thus, the strength of the genomic environment determines how much it will contribute to gene expression, resulting in non-linear scaling of promoter activities across the genome. This is in contrast with the linear scaling we previously observed using a library of proximal enhancers (Maricque et al. 2019), suggesting that core promoters and proximal enhancers may interact with the genomic environment in different ways.

### Genomic clusters have different chromatin states and sequence features

Finally, we asked what features of each cluster distinguish them from each other by overlapping our genomic regions with existing epigenomic datasets and sequence features. Previous studies have shown that reporter genes integrated into the genome tend to take on the chromatin state of the integration site (Chen et al. 2013; Corrales et al. 2017). In general, cluster activity is correlated with chromatin marks associated with active transcription (H3K27ac, H3K4me3) and transcriptional activity (PolII binding, CAGE-seq) (Figs. 6A-C, Supplemental Fig. S10A), while accessible chromatin (ATAC) and CpG methylation do not separate the clusters (Supplemental Figs. S10B and S10C). This suggests that the three clusters are mainly distinguished by their level of transcriptional activity. We also used sequence features to classify the clusters using gapped k-mer SVMs comparing two clusters at a time (Ghandi et al. 2014, 2016). The SVMs performed well, with five-fold cross-validated AUCs ranging from 0.8 to 0.9 (Figs. 6D-F and Supplemental Figs. S7D-F). Scrambling the cluster annotations led to essentially random predictions by the SVM (Supplemental Figs. S7G and S7H). To further validate the model, we used the trained SVM to predict the cluster type of other TRIP integrations that were not in the 5kb region analysis. As expected, clusters that were predicted to be more active also showed higher expression (Supplemental Fig. S7I). To identify the motifs that separate the clusters, we performed *de novo* motif enrichment and identified CG-rich sequences in the more active clusters (Supplemental Figs. S7J and S7K). Similarly, the CG content of each sequence increases from low to high activity clusters on average (Fig. 6G). Motif enrichment using known TF position weight matrices did not identify any obvious enriched TF motifs, suggesting that the clusters are not defined by any single TF. However, when we scanned each sequence for known TF motifs, we find that sequences in more active clusters have more TF motifs than less active clusters on average (Fig. 6H). This result suggests that the differences between clusters is partially explained by the number of TFs binding in each cluster.

**Fig. 6.**
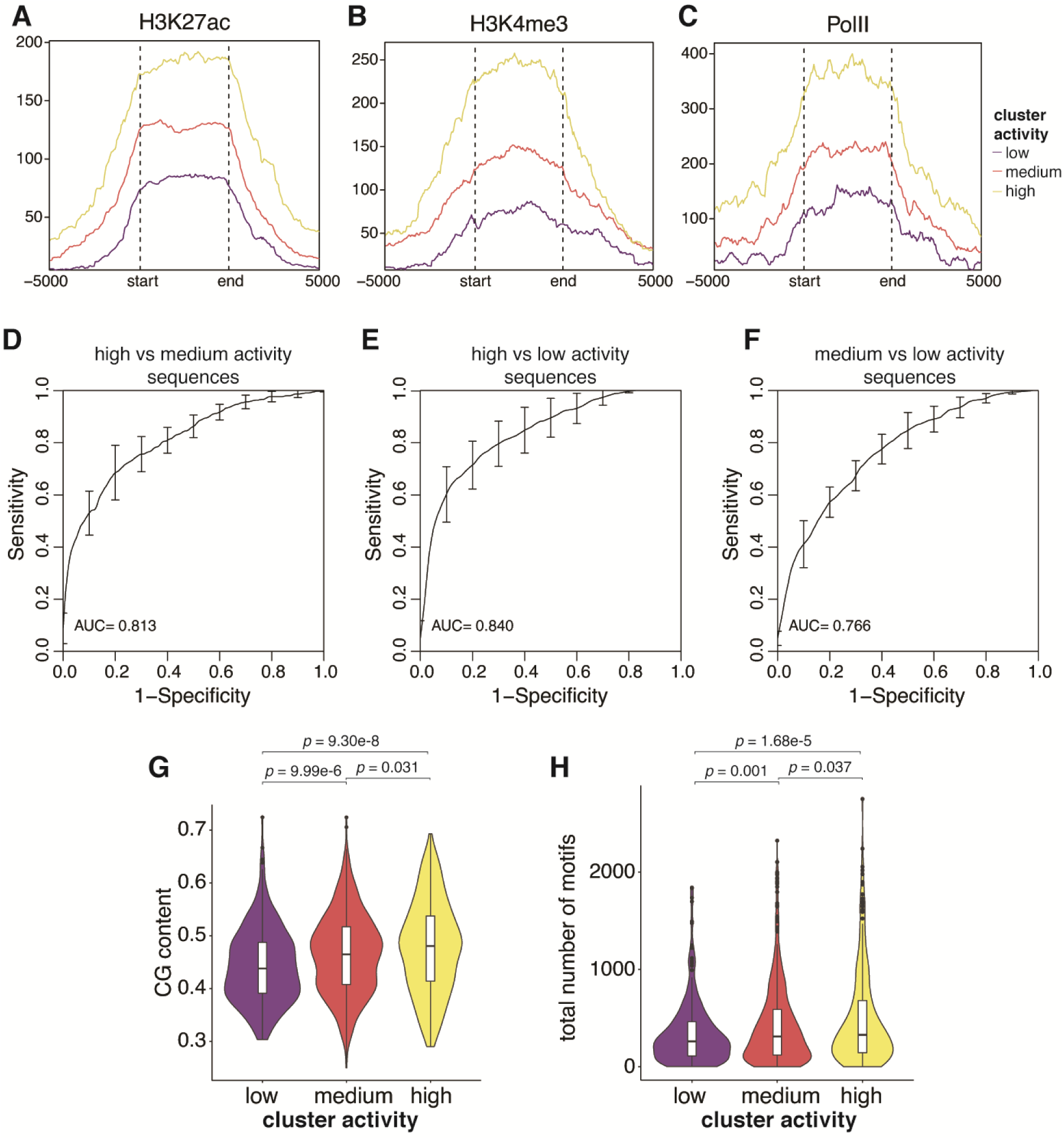
Genomic clusters have different chromatin states and sequence features. **(A, B, C)** Metaplots of H3K27ac, H3K4me3 and PolII levels respectively in each genomic cluster. The start and end marks the boundaries of each genomic region, which are determined by the first and last integration in the region. The x-axis extends +/- 5kb around each genomic region. **(D, E, F)** Performance of gkmSVM used to classify sequences from different genomic clusters. Receiver-operating characteristics (ROC) curves were generated using five-fold cross-validation. **(G)** The GC fraction of each genomic region was calculated and plotted for each cluster. **(H)** Number of TF binding sites in each genomic region was calculated and plotted for each cluster. *p* values were calculated by Student’s *t*-tests.

## Discussion

Gene expression results from the integration of multiple inputs including the core promoter, chromatin environment, distal enhancers and the surrounding transcription factor concentrations. Here we present a framework for dissecting the contributions of core promoters and the surrounding genomic environments to gene expression. Using this framework we found that the intrinsic activities of core promoters are preserved across diverse genomic locations, and are consistent with their activities on plasmids. Contrary to the promoter compatibility hypothesis, hk and dev promoters scale similarly across genomic locations when normalized for differences in strength. These results suggest a general lack of specificity between core promoters and the chromatin landscape/enhancers in their genomic environments, which is consistent with the classical idea of core promoters as passive sequences for the assembly of basal transcriptional machinery. While promoter compatibility has been observed for specific promoter-genomic environment pairs (Ohtsuki et al. 1998; Butler and Kadonaga 2001; Zabidi et al. 2015; Li and Noll 1994; Merli et al. 1996), our results suggest that such interactions are relatively rare or have smaller effects than the effects of genomic scaling. In this model sequence-specific or protein-specific interactions between core promoters and genomic environments contribute less to gene expression than the independent effects of core promoters and genomic environments. This model suggests a modular genome compatible with the evolution of gene expression by genome rearrangements (Carroll 2005; Prud’homme et al. 2007). In a modular genome, core promoters will function in new genomic locations without having to evolve the machinery for a new set of specific interactions at each location.

Unlike our previous results with cis-regulatory sequences upstream of the core promoter, scaling is not a simple linear combination of genomic position effects and promoter effects (Maricque et al. 2019). In a linear relationship, the genomic environment scales the activity of local promoters such that the rank order and quantitative differences between promoters are always preserved. This occurs when the contribution of genomic and promoter effects remains constant across genomic locations and promoters. Instead, we find that the quantitative differences between strong and weak promoters change in different genomic environments (Fig. 5E), suggesting that genomic environments scale core promoter activities in a non-linear manner. We speculate that different core promoter sequence features set the strength of the promoter, which in turn determines how it interacts with the genomic environment. Our data is also consistent with recent simulations showing how promoters starting from different states representing different promoter strengths can have different responses to increasing enhancer contact frequency, giving the appearance of enhancer-promoter specificity (Xiao et al. 2021). In the future, this relationship will allow us to predict gene expression by measuring core promoter strength and genomic environment activity independently.

## Methods

### Library design

We obtained a set of 6916 core promoter sequences from Haberle *et al*. (Haberle et al. 2019) and selected 672 sequences for our library. Each promoter is 133bp long and centered on the major transcription start site (TSS). We selected the sequences to contain diverse core promoter types and expression patterns (Supplemental Table S1 and S2) using the designations obtained from Haberle *et al*.. We also included the super core promoter (SCP1), as well as versions of SCP1 with TATA and DPE single and double mutants (Juven-Gershon et al. 2006).

### patchMPRA library cloning

The core promoter library was synthesized by Agilent technologies through a limited licensing agreement as 200bp oligonucleotides including flanking sequences for cloning. Each element in the library contained 10 barcodes for redundancy, leading to a total of 6760 oligonucleotides. The barcodes were randomly selected from barcode lists generated by the FREE barcodes software (Hawkins et al. 2018). We selected a plasmid with a single enhancer that was shown to drive high expression from the pGL transfer library of our previous patchMPRA experiment (Maricque et al. 2019) as the backbone plasmid. The enhancer contains motifs for Fos/Jun and MAF transcription factors and its full sequence is TGCCCCCCTTCTTCCTATGTCTGATGGAGTTTCCTCTCTAAGTAGCCATTTTATTCTGC TGACTCACCCTCTAACTCCCGGTCTTATTCCATCCTGCCTCAGGGTCTGTGGTGTAGTCATAGCAC. We replaced the hsp68-dsRed construct with the synthesized promoter library including its corresponding BCs and then inserted an mScarlet reporter gene between the promoter and barcodes. To test the library without an upstream enhancer we also cloned the library into a backbone that does not contain the CRS.

### patchMPRA

We replaced the HygTK-GFP cassette in the original landing pad cell lines from Maricque *et al*. (Maricque et al. 2019) with a reporter expressing both TK (thymidine kinase) and BFP. The new cassette contains a functional TK gene, allowing for negative selection of cells that do not have a library member integrated.

K562 cells were maintained in Iscove’s Modified Dulbecco′s Medium (IMDM) + 10% FBS + 1% non-essential amino acids + 1% penicillin/streptomycin. To integrate the library into the genome, we co-transfected the library and CRE recombinase (pBS185 CMV-Cre, Addgene 11916) into 4 K562 ‘landing pad’ cell lines expressing the thymidine kinase (TK) gene (landing pad details in Supplemental Table S3). For each replicate, we transfected 32μg library with 32μg CRE recombinase into 9.6 million total cells using the Neon Transfection System (Life Technologies). We performed 3 separate transfections representing 3 biological replicates. After 3 days, we treated the cells with 2mM ganciclovir to kill the cells that did not successfully integrate a library element. Cells were treated every day for 4 days. We then selected for live cells using the MACS Dead Cell Removal Kit (Miltenyi Biotech) and the cells were allowed to grow until there were sufficient cells for DNA/RNA extractions (about 10 million cells).

DNA and RNA was harvested from the cells using the TRIzol reagent (Life Technologies). The RNA was treated with two rounds of DNase using the Rigorous DNase treatment procedure in the Turbo DNase protocol (Ambion), and cDNA was synthesised with oligo-dT primers using the SuperScript IV First Strand Synthesis System (Invitrogen). The barcodes were then amplified from the cDNA and genomic DNA (gDNA) using the Q5 High Fidelity 2X Master Mix (New England Biolabs) with primers specific to our reporter gene (CPL1-2; Supplemental Table S7). We performed 32 PCRs per cDNA biological replicate and 48 PCRs per gDNA biological replicate, then pooled the PCRs of each replicate for PCR purification. 4ng from each replicate was then further amplified with 2 rounds of PCR to add Illumina sequencing adapters (CPL3-6; Supplemental Table S7). Barcodes were sequenced on the Illumina NextSeq platform.

### Episomal MPRA

We replaced the mScarlet reporter in the patchMPRA library with a tdTomato reporter gene between the promoter and pBC. To ensure that the 3’UTR from the episomal library matches that of the patchMPRA, we further subcloned the library into the landing pad lentiviral vector.

For the MPRA, we transfected the library into K562 cells using the Neon Transfection System (Life Technologies). We performed 2 biological replicates, transfecting 2.4 million cells with 10μg of library per replicate. After 24h, we harvested RNA from the cells using the PureLink RNA Mini Kit (Invitrogen). The RNA was treated with DNase and converted to cDNA in the same way as the patchMPRA library above. We then amplified barcodes from cDNA using primers CPL2 and CPL7 (Supplemental Table S7) with the Q5 High Fidelity 2X Master Mix (New England Biolabs). We performed 4 PCRs per replicate from cDNA. For DNA normalisation, we performed the same PCR (2 PCRs per replicate; 2 replicates) on the plasmid library. The PCRs from the same replicates were then pooled and purified. 4ng from each replicate was then further amplified with 2 rounds of PCR to add Illumina sequencing adapters (CPL3-6; Supplemental Table S7). Barcodes were sequenced on the Illumina NextSeq platform.

### TRIP library cloning

We performed TRIP according to the published protocol with some modifications (Akhtar et al. 2013). Each selected promoter was amplified from the promoter library (CPL8-19; Supplemental Table S7) and cloned into a PiggyBac vector with a unique barcode that identifies the promoter (pBC). Importantly, the promoter and reporter to be integrated is located between two parts of a split-GFP reporter gene (gift from Robi Mitra lab) (Qi et al. 2017). When the promoter-reporter-barcode construct is integrated into the genome, the split-GFP combines to produce functional GFP, allowing us to sort for cells that have successfully integrated the promoters. Each construct was then randomly barcoded by digesting the plasmid with XbaI followed by HiFi assembly (New England Biolabs) with a single-stranded oligo containing 16 random N’s (TRIP barcodes; tBC) and homology arms to the plasmid (CPL20; Supplemental Table S7). Since each promoter is uniquely barcoded, we combined all the promoters into a single library for subsequent TRIP experiments.

### TRIP

The TRIP library and piggyBac transposase (gift from Robi Mitra lab) were co-transfected into wild-type K562 cells at a 1:1 ratio using the Neon Transfection System (Life Technologies). In total, we transfected 4.8 million cells with 16μg each of library and transposase. The cells were sorted after 24 hours for GFP-positive cells to enrich for cells that have integrated the reporters. After a week, the cells were sorted into four pools of 7000 cells each to ensure that each pBC-tBC pair is only integrated once in each pool. The pools were then allowed to grow until there were sufficient cells for DNA/RNA extractions.

We harvested DNA and RNA from the cells using the TRIzol reagent (Life Technologies). The RNA was treated with DNase and converted to cDNA in the same way as the patchMPRA library above. We then amplified barcodes from cDNA and gDNA using primers CPL7 and CPL21 (Supplemental Table S7). We performed 4 PCRs per pool from cDNA and gDNA respectively using the Q5 High Fidelity 2X Master Mix (New England Biolabs), then pooled the PCRs and purified them. 4ng from each replicate was then further amplified with 2 rounds of PCR to add Illumina sequencing adapters (CPL22-23, CPL5-6; Supplemental Table S7). Barcodes were sequenced on the Illumina NextSeq platform.

To map the locations of TRIP integrations, we digested gDNA with a combination of AvrII, NheI, SpeI and XbaI for 16 hours. The digestions were purified and self-ligated at 4°C for another 16 hours. After purifying the ligations, we performed inverse PCR to amplify the barcodes with the associated genomic DNA region (primers CPL24-25; Supplemental Table S7). We did 8 PCRs per pool, purified them and used 4ng of each pool for a further 2 rounds of PCR to add Illumina sequencing adapters (CPL26-28, CPL6; Supplemental Table S7). The library was then sequenced on the Illumina NextSeq platform.

### patchMPRA and episomal MPRA data processing

For patchMPRA, we obtained approximately 11-13 million reads per DNA or RNA replicate from sequencing. For episomal MPRA, we obtained approximately 500,000 reads per DNA or RNA replicate. Reads that contained the barcodes in the proper sequence context were included in subsequent analysis. The pBCs were then decoded using the FREE barcodes software (Hawkins et al. 2018) and the expression of each barcode pair was calculated as log_2_(RNA/DNA). We averaged the expression of barcodes corresponding to the same promoter within each replicate to get promoter expression per replicate, then averaged across replicates for subsequent downstream analysis. Expression values can be found in Supplemental Table S8 (patchMPRA) and Supplemental Table S9 (episomal MPRA).

### TRIP data processing

We obtained approximately 14-25 million reads per DNA or RNA pool from sequencing. Reads that contained both the tBC and pBC in the proper sequence context were included in subsequent analysis. We further filtered tBCs such that they are at least 3 hamming distance apart from every other barcode to account for mutations that occurred during PCR and sequencing. The expression of each BC pair was calculated as log_2_(RNA/DNA). We added a pseudocount to the RNA counts to include barcode pairs that had DNA but no RNA reads. Data from the 4 independent pools were combined in all analyses. Expression values can be found in Supplemental Table S5.

For the locations of TRIP integrations, reads containing each barcode pair were matched with the sequence of its integration site. The integration site sequences were then aligned to hg38 using *bwa* with default parameters. Only barcodes that mapped to a unique location were kept for downstream analyses. The mapped integration locations can be found in Supplemental Table S5.

### TRIP data analysis

We downloaded a list of expressed genes in K562 cells using whole cell long polyA RNA-seq data generated by ENCODE (Djebali et al. 2012) from the EMBL Expression Atlas. We then designated the genes as hk or dev based on the list of hk genes obtained from Eisenberg and Levanon (Eisenberg and Levanon 2013). Using the locations of these promoters (GENCODE Release 36, GRCh38.p13) we identified TRIP integrations located within 5kb of either hk or dev promoters and plotted the expression of these integrations separately.

To increase the resolution of the analysis we identified genomic regions where at least 4 different promoters integrated within 5kb of each other (Full list of regions in Supplemental Table S6). For regions in which the same promoter integrated more than once we used the median expression of that promoter. This yielded 1268 genomic regions. All heatmaps were generated using the ComplexHeatmap package in R (Gu et al. 2016). To determine the diversity of the identified 5kb regions, we downloaded the 15-state segmentation for K562 (hg19) from the ENCODE portal and performed a liftover to hg38 using the UCSC liftover tool (Hinrichs et al. 2006). We then overlapped the 5kb regions with chromHMM regions using a minimum overlap of 200bp using the Genomic Ranges R packages (Lawrence et al. 2013).

To rank and cluster the regions we first imputed missing values using the mean of the promoter across all locations. We then used the means of each region to rank the clusters and plotted the smoothed expression of each promoter. To cluster the 5kb genomic regions, we ran k-means clustering on the imputed data using the ConsensusClusterPlus package in R (Wilkerson and Hayes 2010). The imputed data was only used for ranking and clustering and not downstream analysis.

### Epigenome data analysis

For the cluster metaplots, we considered the boundaries of each genomic region as the locations of the first and last integrations in each region. We then downloaded various K562 epigenome datasets (full list of sources in Supplemental Table S10). For CpG methylation, we downloaded both replicates and used the averaged signal from both replicates. For H3K27ac, H3K4me3, PolII, CpG methylation and ATAC-seq, we used the EnrichedHeatmap package in R (Gu et al. 2018) to draw the metaplots for each cluster extending 5kb upstream and downstream of each genomic region. For CAGE-seq, we downloaded the hg19 dataset from the FANTOM5 consortium (Lizio et al. 2015, 2019) and converted it to hg38 using the UCSC liftover tool (Hinrichs et al. 2006). Because the signal was relatively sparse across genomic locations, we plotted the total CAGE signal across each genomic region.

### Sequence features analysis

We obtained the sequences of each region using the BSgenome package in R (Pagès 2020). For the gapped k-mer predictions, we used the gkmSVM R package (Ghandi et al. 2016) with word length = 10 and number of informative columns = 6. We used AME for motif enrichment analysis (McLeay and Bailey 2010), DREME for *de novo* motif discovery (Bailey 2011) and FIMO to determine the number of motifs per sequence (Grant et al. 2011), from MEME suite 5.0.4. For all motif analyses we limited analysis to expressed transcription factors (FPKM >= 1) in K562 from whole cell long polyA RNA-seq data generated by ENCODE (Djebali et al. 2012) downloaded from the EMBL Expression Atlas.

To predict the type of genomic region of other integrations not in the defined 5kb regions, we obtained genomic sequences of the 1kb flanking region around the integration (500bp upstream and 500bp downstream). We then used the trained gkmSVM kernels to calculate the weights of each flanking region and assigned the integrations into low, medium or high activity clusters based on their weights. Only integrations that could be confidently assigned were included.

### Modeling

We fit log_2_ expression values with linear models of core promoter and genomic location activities using the lm function in R. Variance explained by each term was calculated with one-way ANOVAs of the respective models.

## Data Access

All raw and processed sequencing data generated in this study have been submitted to the NCBI Gene Expression Omnibus (GEO; https://www.ncbi.nlm.nih.gov/geo/) under accession number GSE173678. The code used to process the data can be found at https://github.com/claricehong/core_promoters_2021.

## Acknowledgments

We are grateful to Vanja Haberle and Alexander Stark for providing us with sequences from their core promoter library and helpful suggestions for which promoters to select. We also thank Ting Wang, Brett Maricque and members of the Cohen Lab for their helpful comments and critical feedback on the manuscript; Jessica Hoisington-Lopez and MariaLynn Crosby in the DNA Sequencing Innovation Lab for assistance with high-throughput sequencing and the Genome Engineering and iPSC Center for kindly allowing us to use their flow cytometer for cell sorting. This work was supported by grants to BAC from the National Institutes of Health (R01GM092910).

---

**Supplemental Fig. S1.**
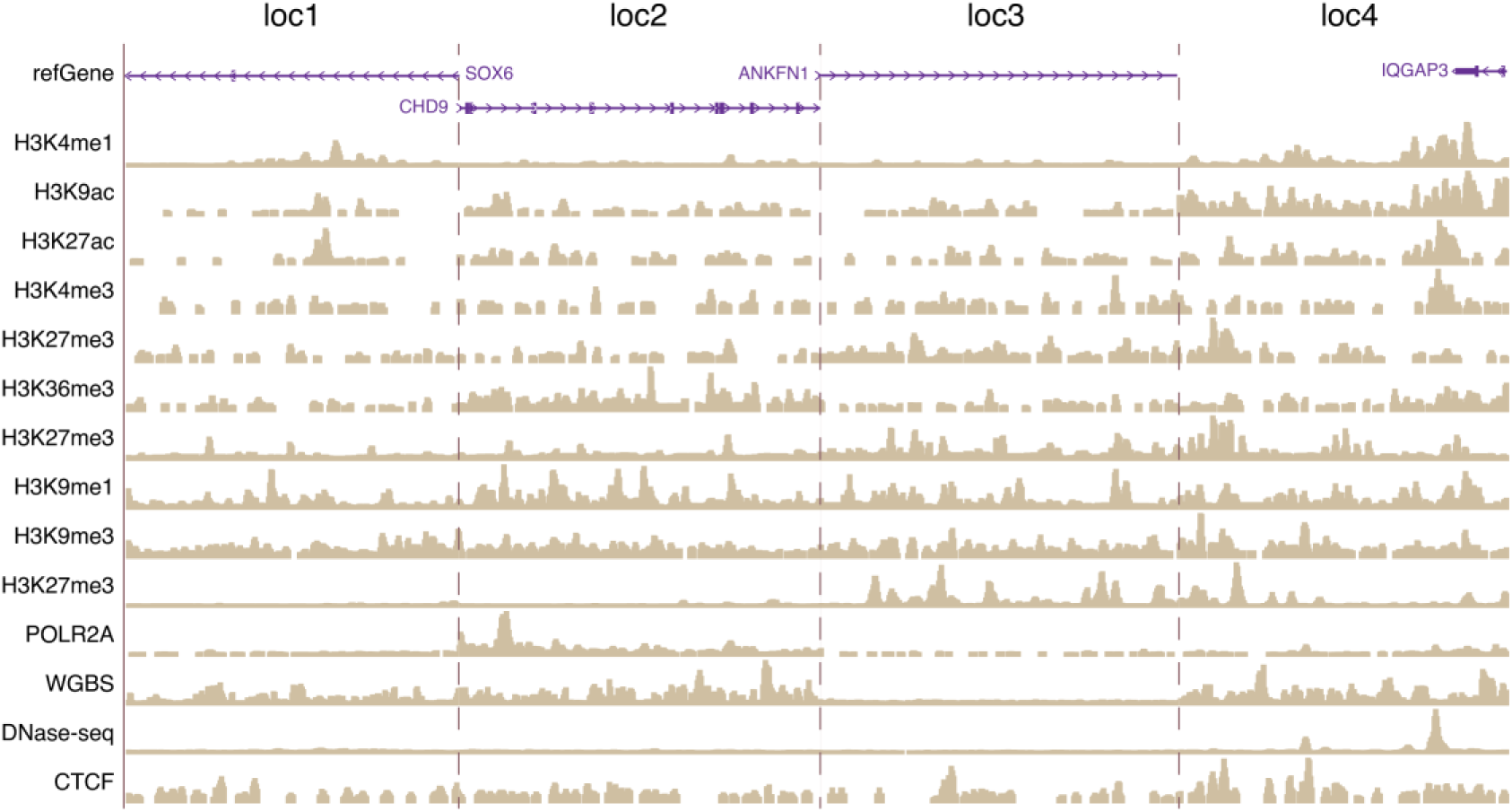
Landing pad locations have diverse chromatin marks and transcriptional activity. Each section is centered on the location of landing pad integration.

**Supplemental Fig. S2.**
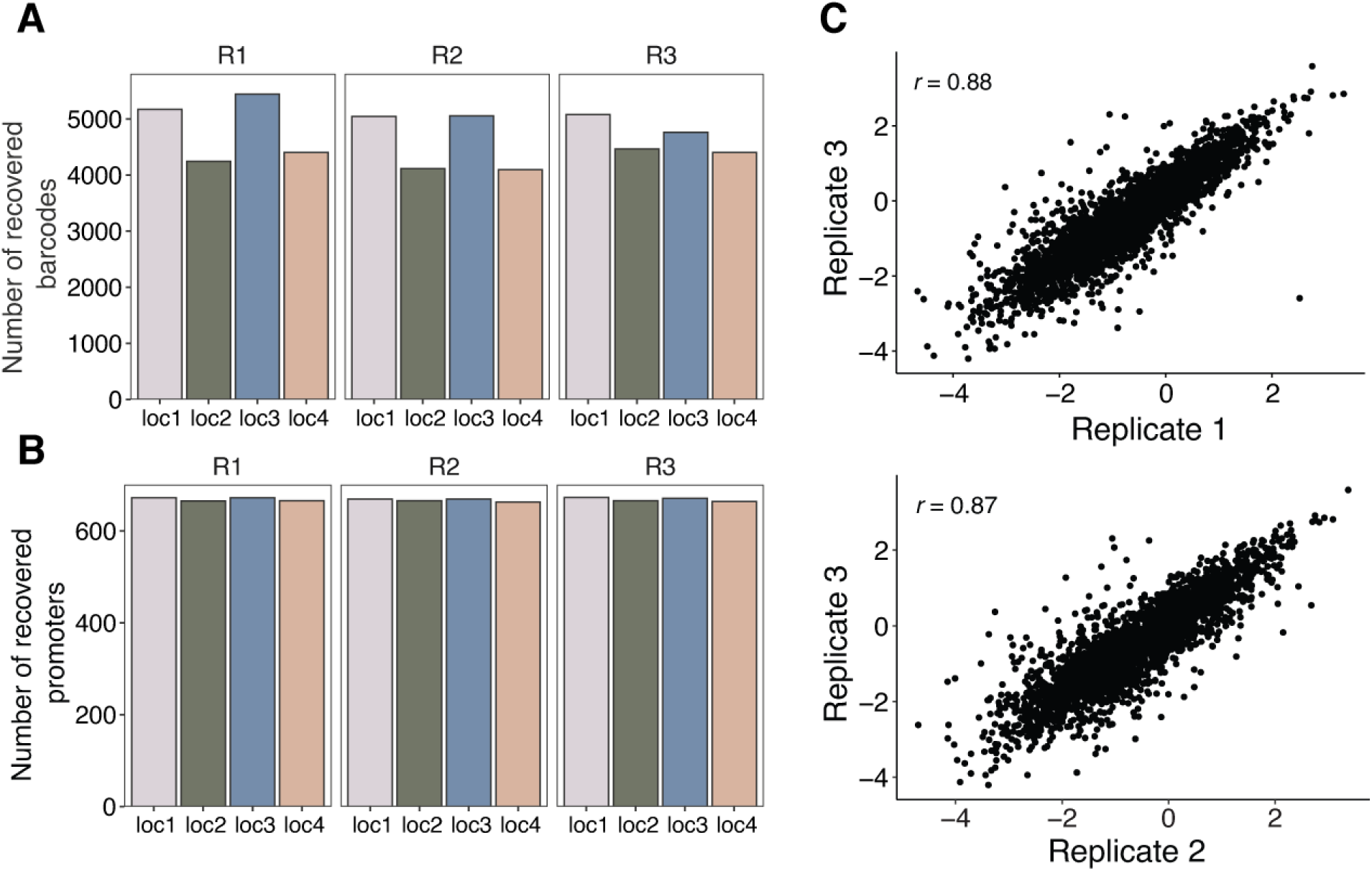
patchMPRA measurements are reproducible. **(A)** Number of promoter barcodes recovered from each location per biological replicate. **(B)** Number of promoters recovered from each location per biological replicate. **(C)** Reproducibility of core promoter measurements from independent patchMPRA transfections.

**Supplemental Fig. S3.**
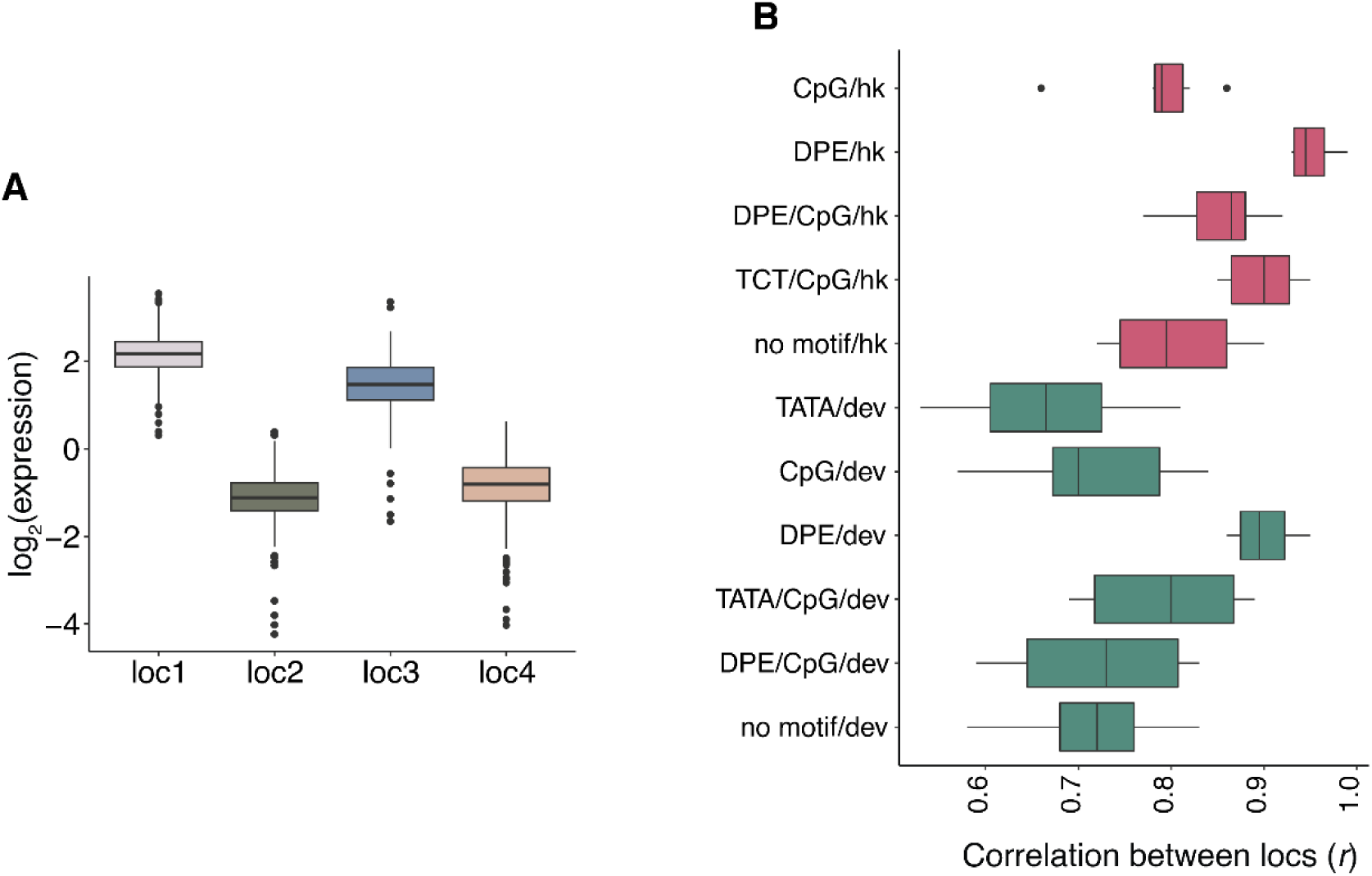
Effect of genomic locations on core promoters. **(A)** Expression of a library of proximal enhancers at each genomic location (Maricque et al. 2019). **(B)** All pairwise correlations (Pearson’s *r*) between genomic locations for core promoters with different motifs within each class.

**Supplemental Fig. S4.**
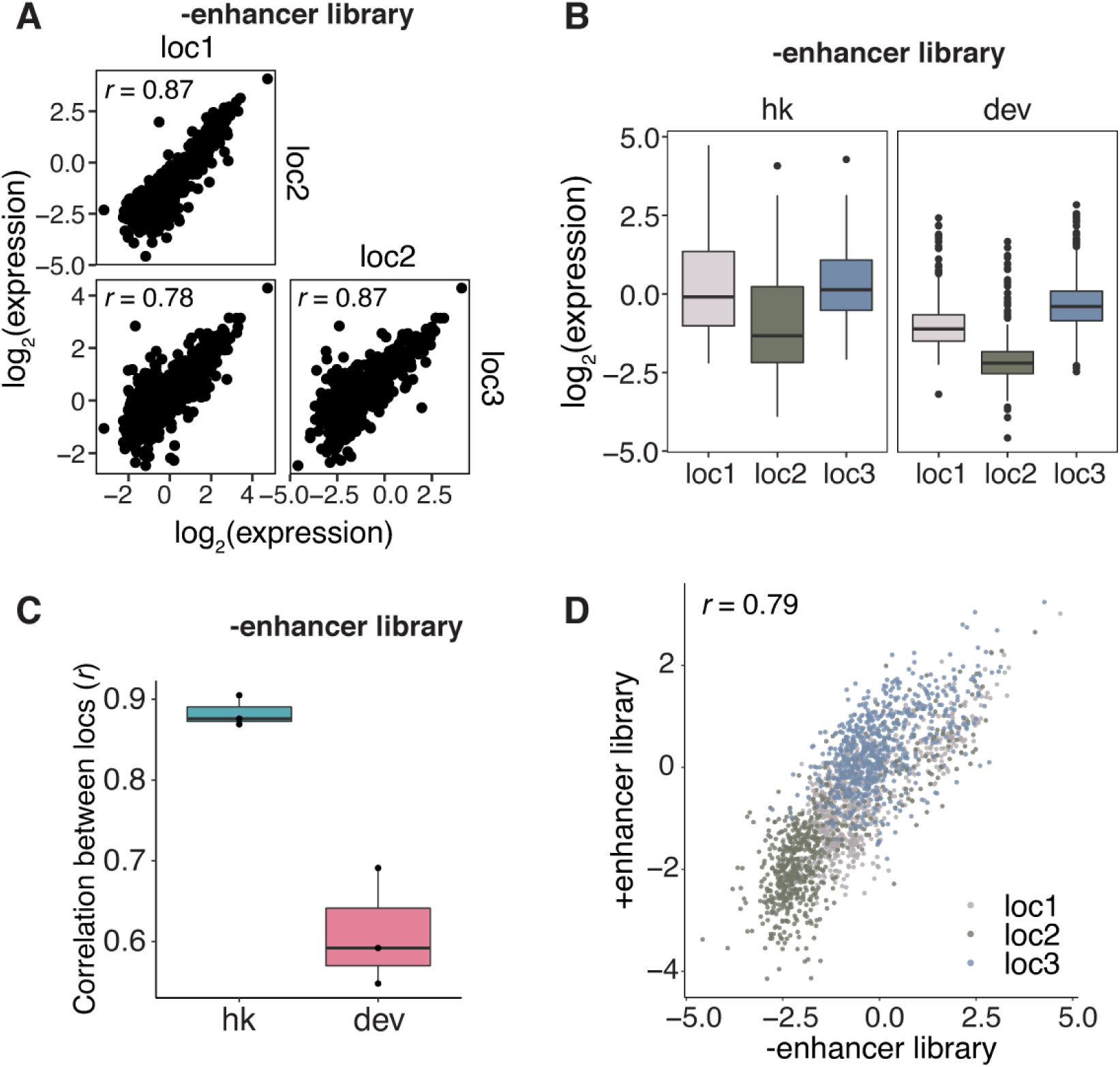
Core promoter scaling is not dependent on the proximal enhancer. **(A)** Pairwise correlations (Pearson’s *r*) of core promoter activity between the different genomic locations after removing the proximal enhancer upstream of the core promoter. **(B)** Expression of hk and dev promoters at each genomic location without the proximal enhancer. **(C)** All pairwise correlations (Pearson’s *r*) between genomic locations for hk and dev core promoters without the proximal enhancer. **(D)** Correlation between expression with and without the proximal enhancer at locations 1-3.

**Supplemental Fig. S5.**
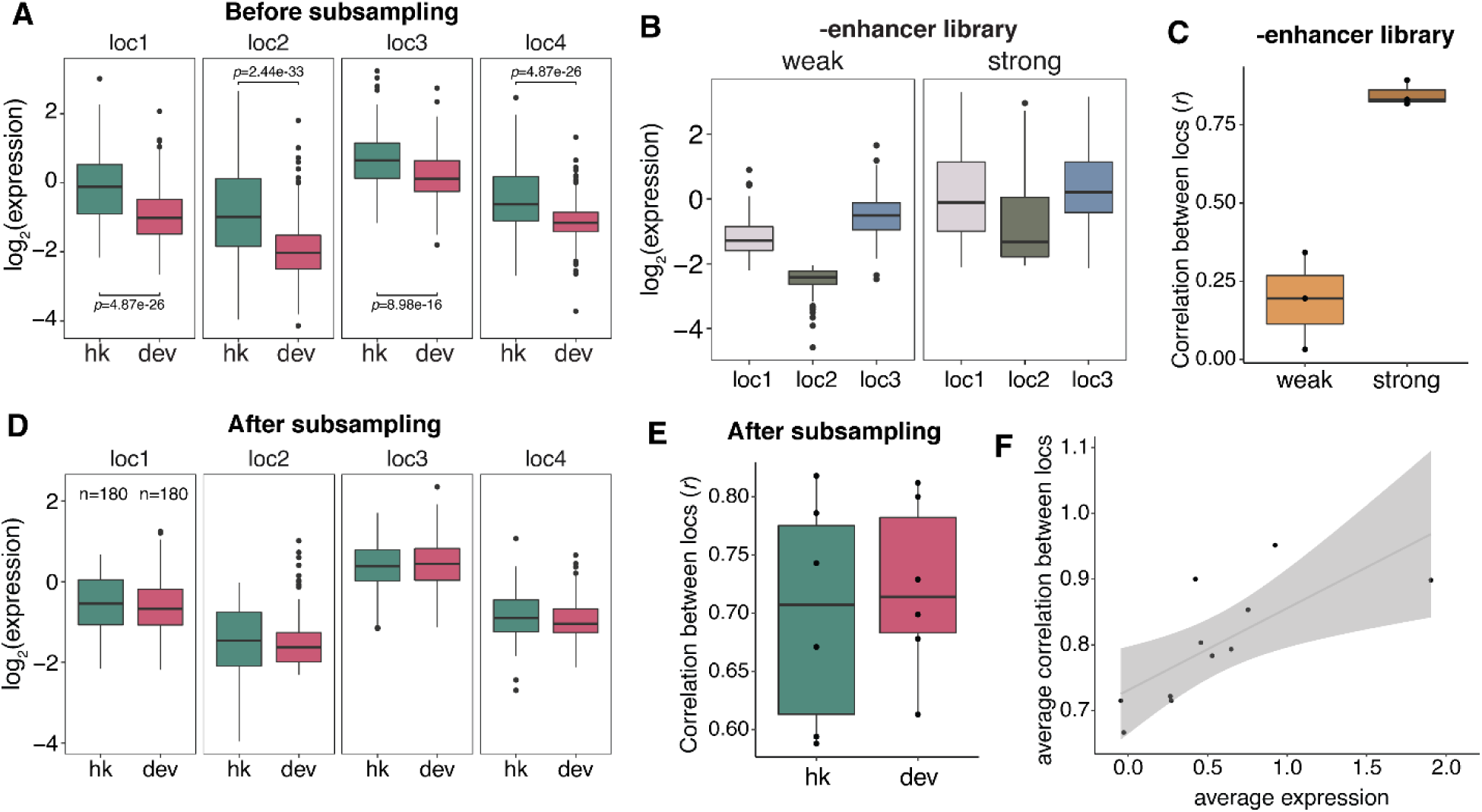
Intrinsic promoter strength explains differences between classes of core promoters. **(A)** Expression of all hk and dev promoters at each genomic location. *p*-values were calculated by Student’s *t*-tests. **(B)** The effect of genomic location on the expression of weak and strong core promoters after removing the proximal enhancer upstream of the promoter. **(C)** All pairwise correlations between genomic locations for weak and strong core promoters without the proximal enhancer. **(D)** Expression of hk and dev promoters at each genomic location after sampling promoters such that the two classes have equivalent average strengths. n indicates number of promoters sampled from each class. **(E)** All pairwise correlations between genomic locations for subsampled hk and dev core promoters. **(F)** The pairwise correlations of core promoters with different motifs (from Supplemental Fig. S2) are explained by the average expression of each group.

**Supplemental Fig. S6.**
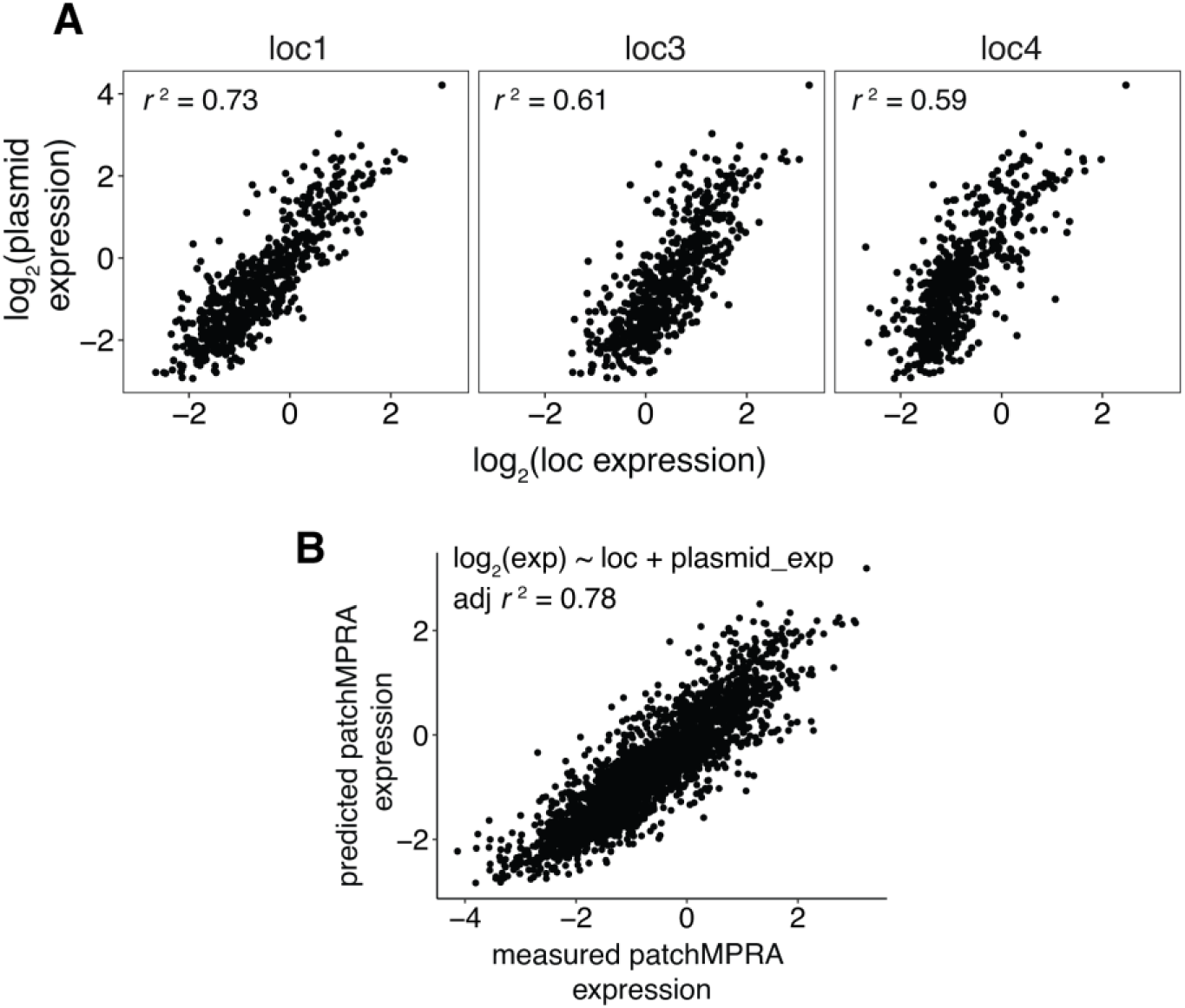
Core promoter activities in the genome reflect the promoters’ intrinsic activity. **(A)** Correlations between expression of core promoter library measured on plasmids and at the indicated genomic location by patchMPRA. **(B)** Correlation between measured expression by patchMPRA and predicted expression by a linear model using core promoter intrinsic activity measured on plasmids. All correlations were calculated as Pearson’s *r*.

**Supplemental Fig. S7.**
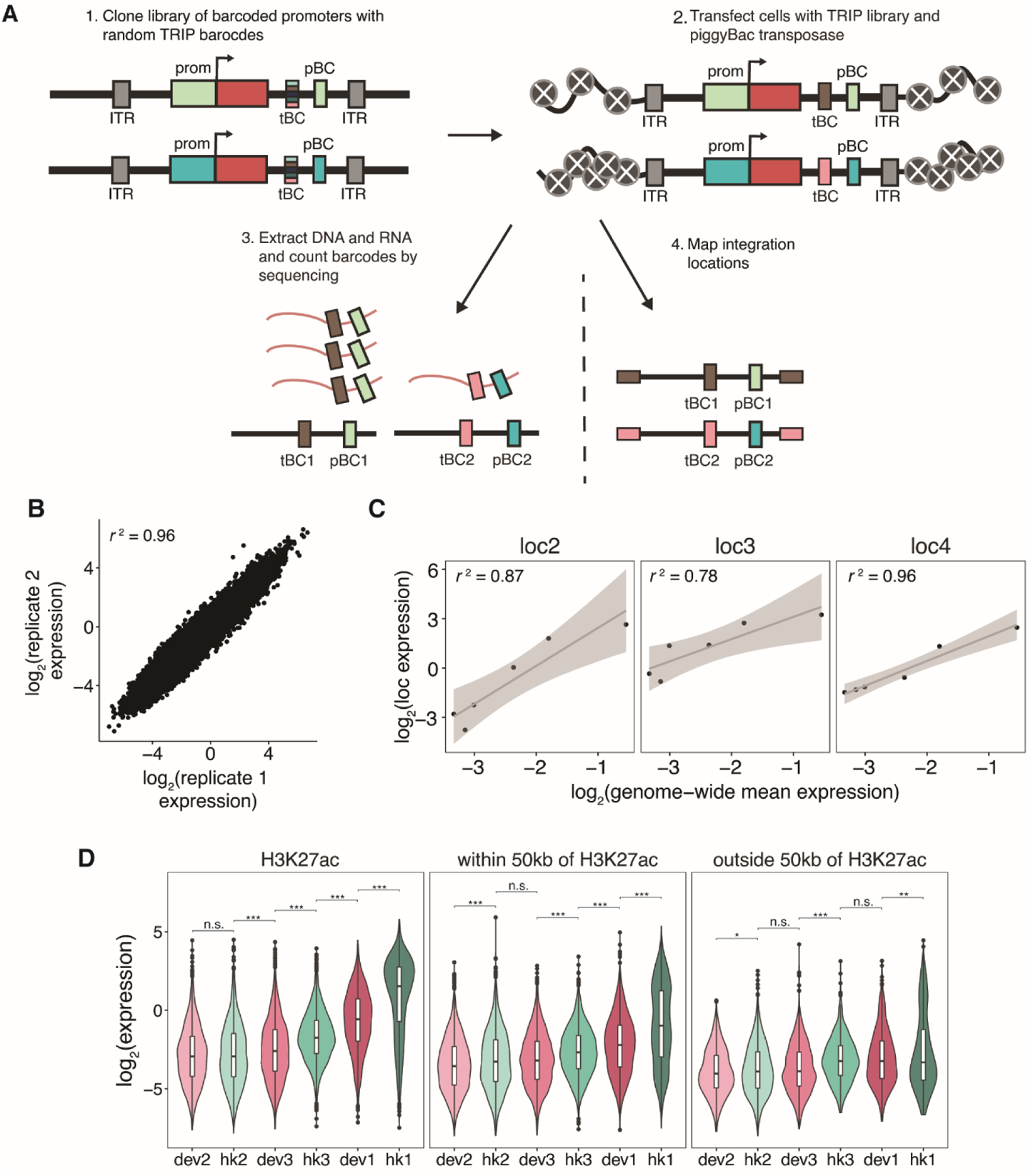
Measurements of six core promoters at thousands of genomic locations by TRIP. **(A)** Schematic of TRIP experiment. tBC: TRIP barcode; pBC: promoter barcode; ITR: inverted terminal repeats. **(B)** Reproducibility between measurements from independent DNA and RNA extractions. **(C)** Correlations between mean expression of core promoters measured by TRIP and at the indicated genomic location by patchMPRA. **(D)** Expression of core promoters at locations that overlap with H3K27ac sites, are within 50kb of H3K27ac sites or are outside 50kb of H3K27ac sites. *p*-values were calculated by Student’s *t*-tests. n.s.: not significant, * *p*<0.05, ** *p*<0.001, *** *p*<0.005.

**Supplemental Fig. S8.**
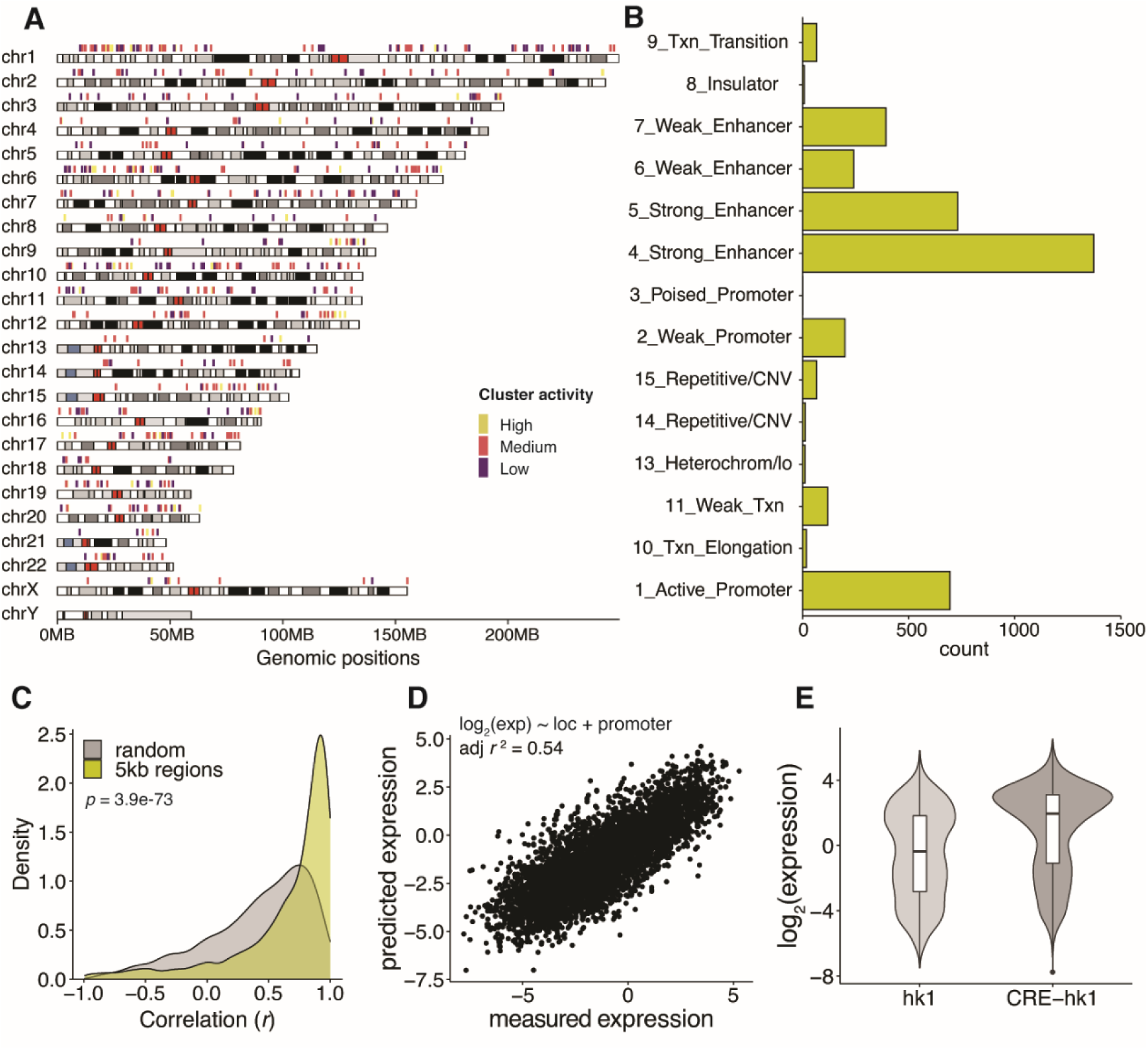
Core promoter scaling is a genome-wide phenomenon. **(A)** Regions with ≥4 different promoters integrated within 5kb of each other are located across the genome. Cluster activity was designated by the analysis in Fig. 5B. **(B)** Distribution of chromHMM annotations of defined 5kb regions. **(C)** For each defined 5kb region, correlations (Pearson’s *r*) between core promoter activity measured by TRIP and by patchMPRA were calculated and all correlations were plotted as a density plot. As a comparison, we randomly grouped promoters without considering their integration locations and calculated the correlations for each group. The *p* value was calculated using the Mann–Whitney *U* test. **(D)** Correlation between measured expression by TRIP and predicted expression using a model assuming independence between genomic environments and core promoters. **(E)** Expression of all integrations of hk1 and hk1 with an upstream *cis*-regulatory enhancer (CRE-hk1).

**Supplemental Fig. S9.**
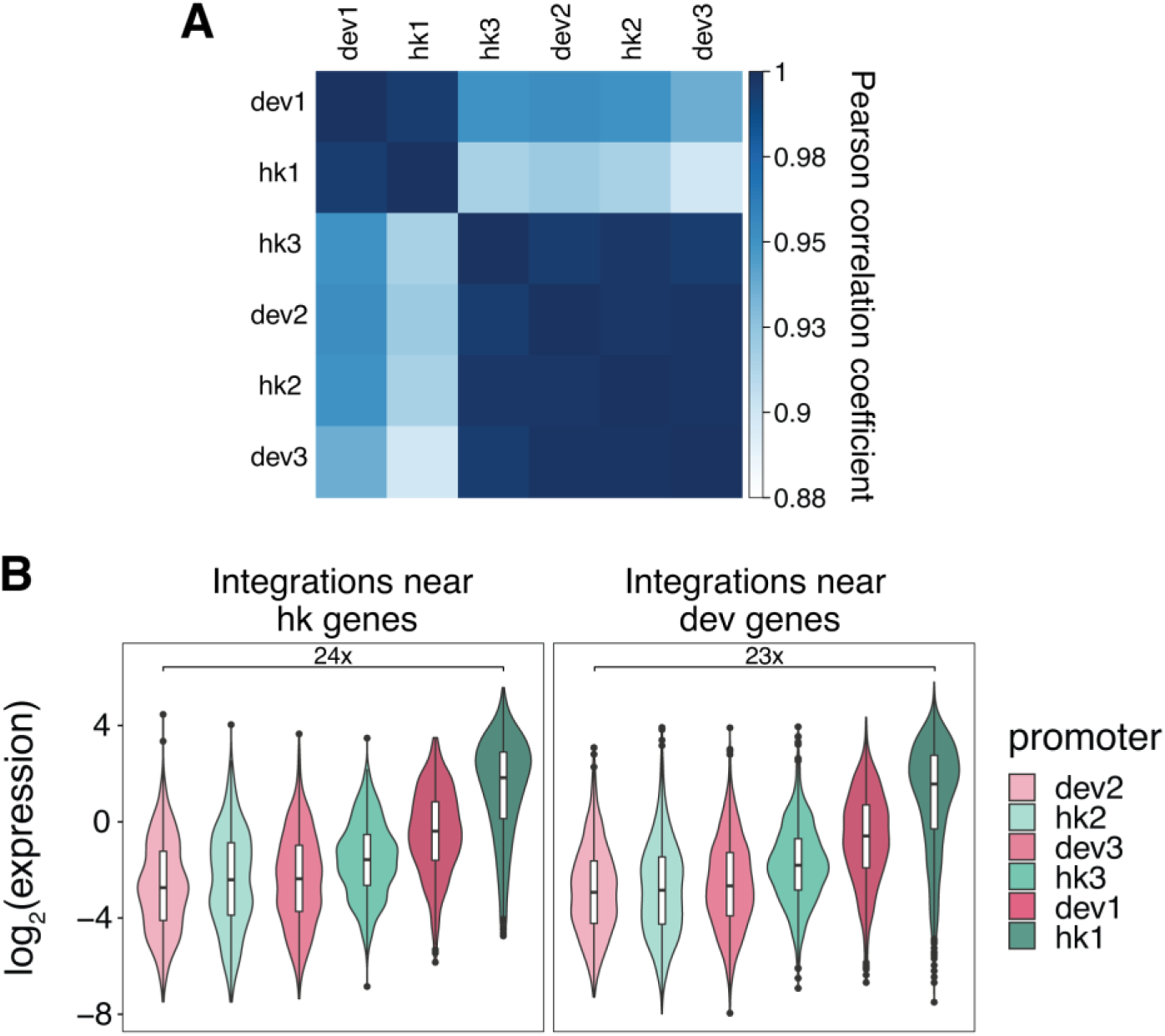
Promoter strength, not class, determines its interaction with the genomic environment. **(A)** Correlation coefficients between curves fitted on each promoter in Fig. 5A. **(B)** Hk and dev integrated core promoters behave similarly near endogenous hk or dev promoters.

**Supplemental Fig. S10.**
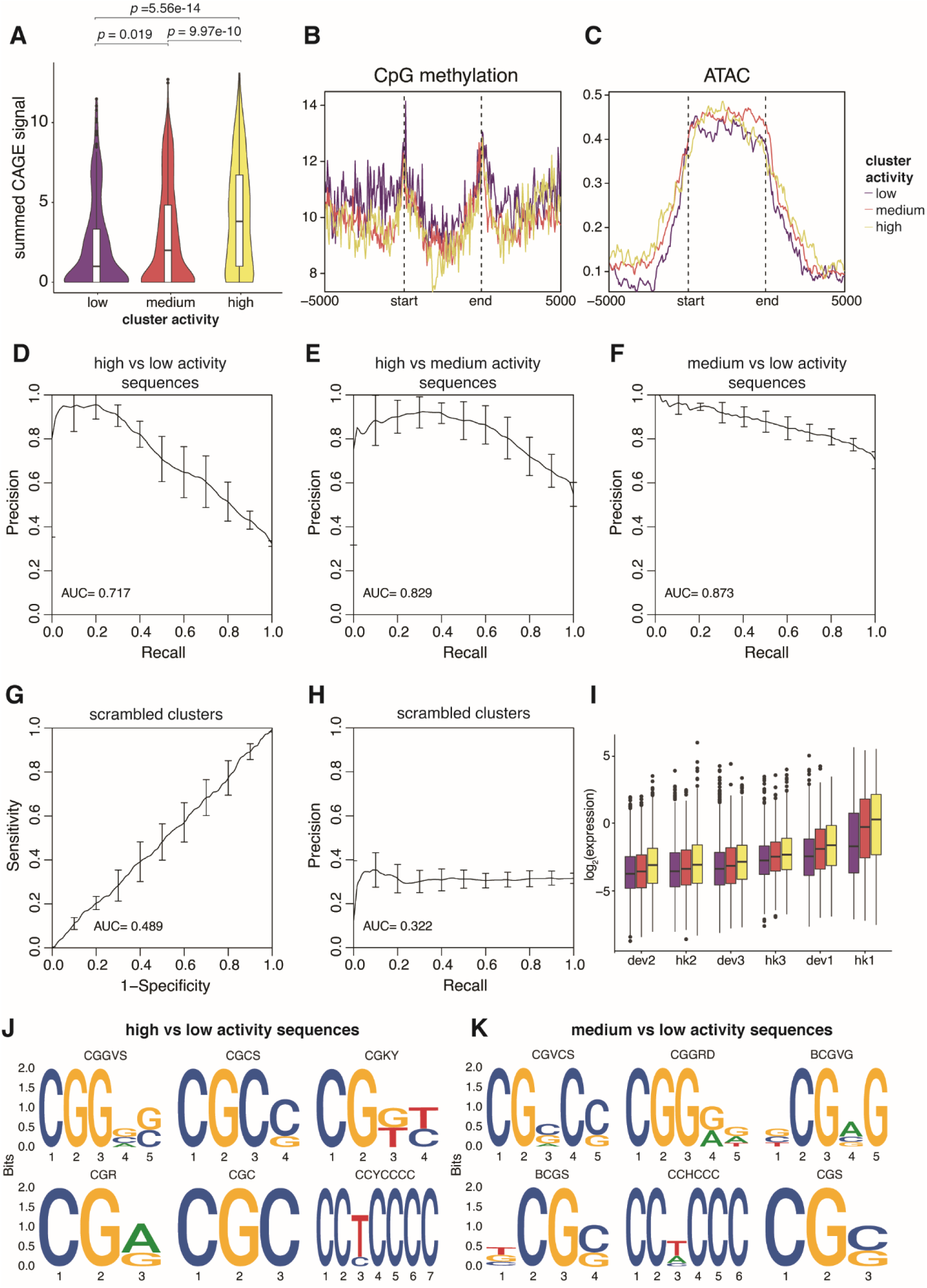
Epigenomic signatures and sequence features of different genomic activity clusters. **(A**) CAGE-seq signal was calculated for each genomic region, and the summed signals were plotted for each cluster. *p* values were calculated by Student’s *t*-tests. **(B, C)** Metaplots of CpG methylation and ATAC-seq signals respectively in each genomic cluster. The start and end mark the boundaries of each genomic region, which are determined by the first and last integration in the region. The x-axis extends +/- 5kb around each genomic region. **(D, E, F)** Performance of gkmSVM used to classify sequences from different genomic clusters. Precision-recall curves (PRCs) were generated using five-fold cross-validation. **(G, H)** Performance of gkmSVM on sequences with scrambled cluster assignments. **(I)** TRIP integrations that were not included in the 5kb genomic region analysis were assigned to a cluster based on their sequence features from the gkmSVM, and the expression of each promoter was plotted based on their predicted clusters. **(J, K)** Top 6 motifs identified by *de novo* motif finding comparing high/low and medium/low activity sequences respectively.

**Supplemental Table S1.** Composition of promoter classes in the core promoter library.

| Promoter class | hk/dev | Number |
| --- | --- | --- |
| TATA-box | dev | 100 |
| CpG island | hk | 100 |
| CpG island | dev | 100 |
| TCT | hk | 2 |
| DPE | hk | 8 |
| DPE | dev | 13 |
| No known motif | hk | 100 |
| No known motif | dev | 100 |
| TATA-box & CpG island | dev | 63 |
| TATA-box & DPE | dev | 2 |
| DPE & CpG island | dev | 14 |
| DPE & CpG island | hk | 43 |
| TCT & CpG island | hk | 22 |
| SCP (and mutants) | - | 4 |
| Total |  | 676 |

**Supplemental Table S3.** Locations of four landing pads in patchMPRA.

| **Name** | **Chr** | **Location (hg19)** | **Annotation** | **Name in Maricque *et al.* (29)** |
| --- | --- | --- | --- | --- |
| loc1 | chr11 | 16,258,750 | Sox6 Intron | LP3 |
| loc2 | chr16 | 53,275,015 | CHD9 Intron | LP4 |
| loc3 | chr17 | 56,426,171 | Intergenic | LP5 |
| loc4 | chr1 | 156,489,766 | Intergenic | LP6 |

**Supplemental Table S7.**
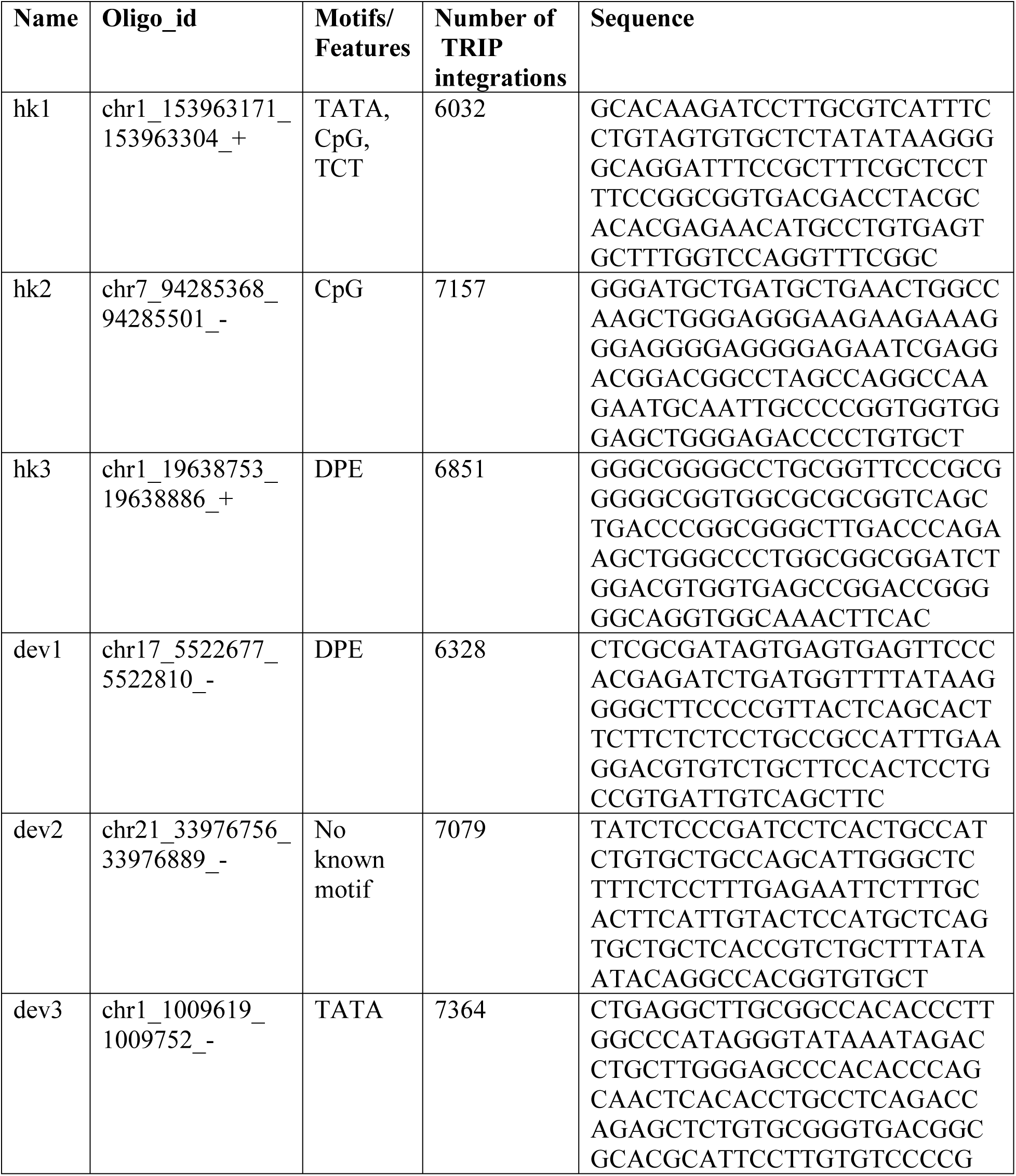
Promoters selected for TRIP.

**Supplemental Table S7.**
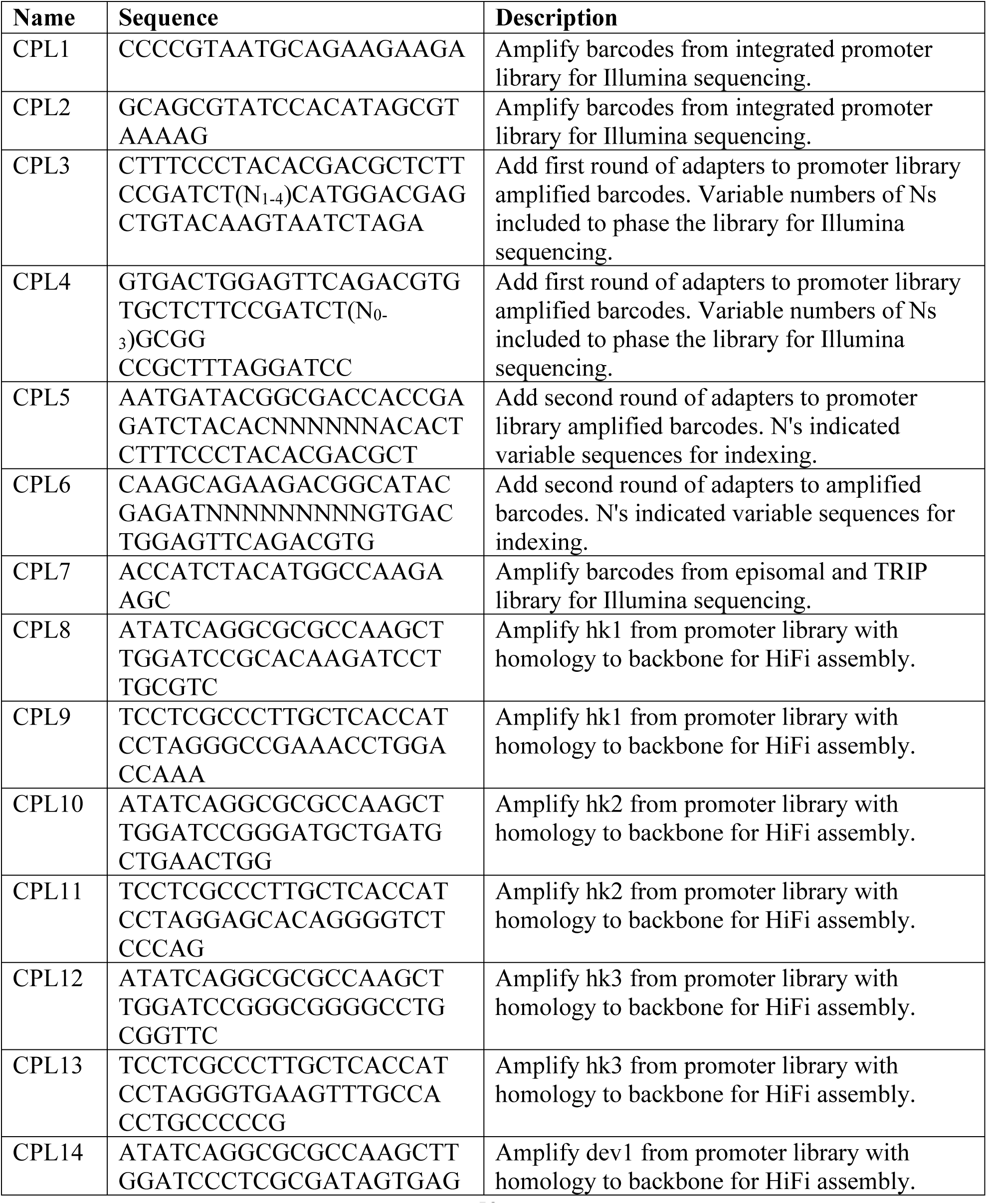
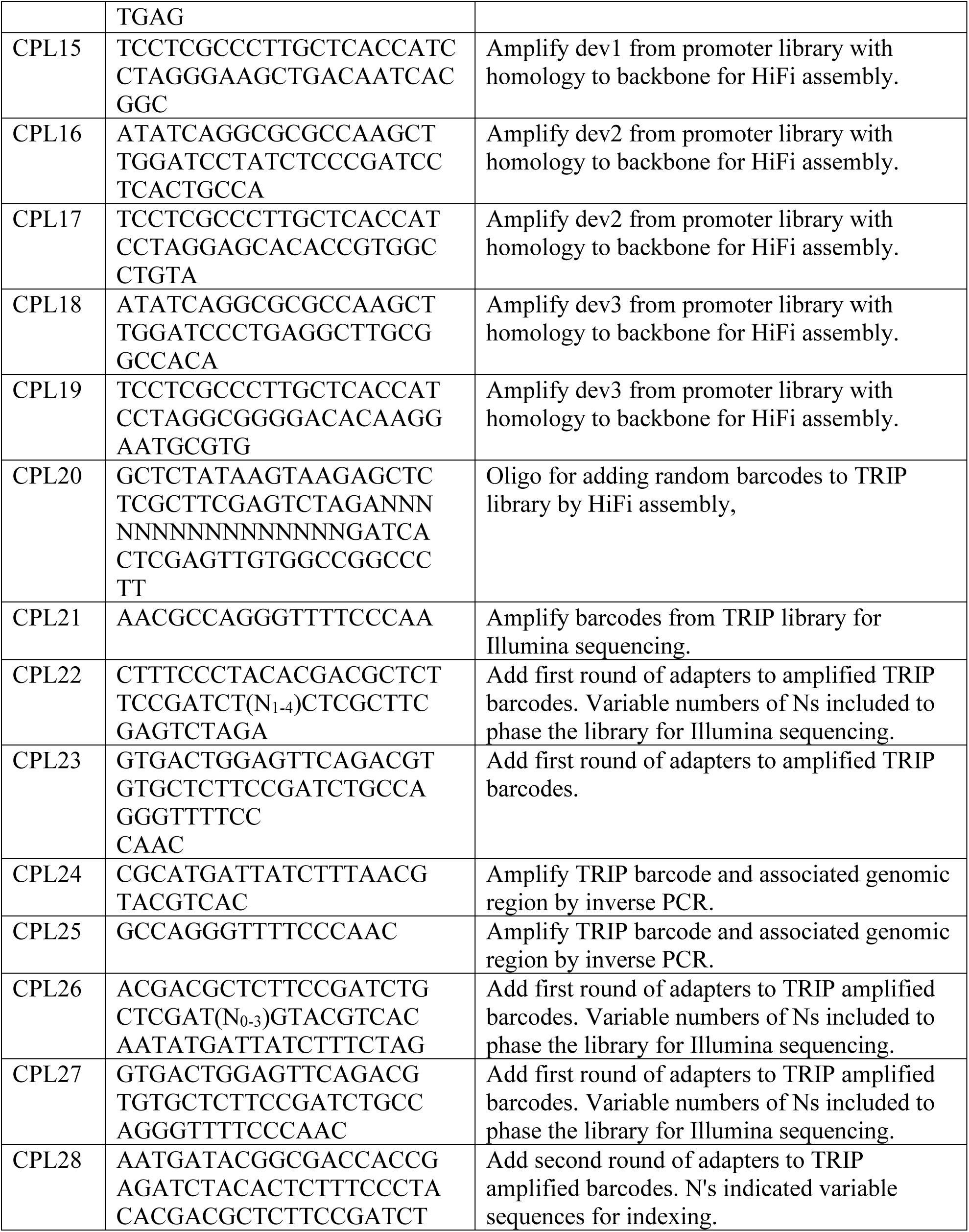
Primers used in this study.

**Supplemental Table S10.** Sources of epigenome datasets used in this study.

| Data | Source Experiment | Source File |
| --- | --- | --- |
| H3K27ac<br>ChIP-seq | ENCSR000AKP | ENCFF437DPT |
| H3K4me3<br>ChIP-seq | ENCSR000EWA | ENCFF916MPM |
| PolII ChIP-seq | ENCSR388QZF | ENCFF285MBX |
| CAGE | FANTOM5 | chronic%20myelogenous%20leukemia%20cell%20line%3aK562.CNhs11250.10454106G4.hg19.ctss.bed.gz |
| CpG<br>methylation | ENCSR765JPC | ENCFF867JRG; ENCFF721JMB |
| ATAC-seq | ENCSR868FGK | ENCFF698MIQ |

## References

Akhtar W, de Jong J, Pindyurin AV, Pagie L, Meuleman W, de Ridder J, Berns A, Wessels LFA, van Lohuizen M, van Steensel B. 2013. Chromatin Position Effects Assayed by Thousands of Reporters Integrated in Parallel. Cell 154: 914–927.

Arnold CD, Zabidi MA, Pagani M, Rath M, Schernhuber K, Kazmar T, Stark A. 2017. Genome-wide assessment of sequence-intrinsic enhancer responsiveness at single-base-pair resolution. Nat Biotechnol 35: 136–144.

Bailey TL. 2011. DREME: motif discovery in transcription factor ChIP-seq data. Bioinformatics 27: 1653–1659.

Butler JEF, Kadonaga JT. 2001. Enhancer–promoter specificity mediated by DPE or TATA core promoter motifs. Genes Dev 15: 2515–2519.

Carroll SB. 2005. Evolution at Two Levels: On Genes and Form. PLOS Biol 3: e245.

Chen M, Licon K, Otsuka R, Pillus L, Ideker T. 2013. Decoupling Epigenetic and Genetic Effects through Systematic Analysis of Gene Position. Cell Rep 3: 128–137.

Corrales M, Rosado A, Cortini R, van Arensbergen J, van Steensel B, Filion GJ. 2017. Clustering of Drosophila housekeeping promoters facilitates their expression. Genome Res 27: 1153–1161.

Djebali S, Davis CA, Merkel A, Dobin A, Lassmann T, Mortazavi A, Tanzer A, Lagarde J, Lin W, Schlesinger F, et al. 2012. Landscape of transcription in human cells. Nature 489: 101–108.

Eisenberg E, Levanon EY. 2013. Human housekeeping genes, revisited. Trends Genet 29: 569–574.

Emami KH, Navarre WW, Smale ST. 1995. Core promoter specificities of the Sp1 and VP16 transcriptional activation domains. Mol Cell Biol 15: 5906–5916.

Ernst J, Kellis M. 2010. Discovery and characterization of chromatin states for systematic annotation of the human genome. Nat Biotechnol 28: 817–825.

Ernst J, Kheradpour P, Mikkelsen TS, Shoresh N, Ward LD, Epstein CB, Zhang X, Wang L, Issner R, Coyne M, et al. 2011. Mapping and analysis of chromatin state dynamics in nine human cell types. Nature 473: 43–49.

Gehrig J, Reischl M, Kalmár É, Ferg M, Hadzhiev Y, Zaucker A, Song C, Schindler S, Liebel U, Müller F. 2009. Automated high-throughput mapping of promoter-enhancer interactions in zebrafish embryos. Nat Methods 6: 911–916.

Ghandi M, Lee D, Mohammad-Noori M, Beer MA. 2014. Enhanced regulatory sequence prediction using gapped k-mer features. PLoS Comput Biol 10: e1003711.

Ghandi M, Mohammad-Noori M, Ghareghani N, Lee D, Garraway L, Beer MA. 2016. gkmSVM: an R package for gapped-kmer SVM. Bioinforma Oxf Engl 32: 2205–2207.

Grant CE, Bailey TL, Noble WS. 2011. FIMO: scanning for occurrences of a given motif. Bioinformatics 27: 1017–1018.

Gu Z, Eils R, Schlesner M. 2016. Complex heatmaps reveal patterns and correlations in multidimensional genomic data. Bioinforma Oxf Engl 32: 2847–2849.

Gu Z, Eils R, Schlesner M, Ishaque N. 2018. EnrichedHeatmap: an R/Bioconductor package for comprehensive visualization of genomic signal associations. BMC Genomics 19: 234.

Haberle V, Arnold CD, Pagani M, Rath M, Schernhuber K, Stark A. 2019. Transcriptional cofactors display specificity for distinct types of core promoters. Nature 570: 122–126.

Haberle V, Stark A. 2018. Eukaryotic core promoters and the functional basis of transcription initiation. Nat Rev Mol Cell Biol 19: 621–637.

Hawkins JA, Jones SK, Finkelstein IJ, Press WH. 2018. Indel-correcting DNA barcodes for high-throughput sequencing. Proc Natl Acad Sci 115: E6217–E6226.

Hinrichs AS, Karolchik D, Baertsch R, Barber GP, Bejerano G, Clawson H, Diekhans M, Furey TS, Harte RA, Hsu F, et al. 2006. The UCSC Genome Browser Database: update 2006. Nucleic Acids Res 34: D590–598.

Juven-Gershon T, Cheng S, Kadonaga JT. 2006. Rational design of a super core promoter that enhances gene expression. Nat Methods 3: 917–922.

Lawrence M, Huber W, Pagès H, Aboyoun P, Carlson M, Gentleman R, Morgan MT, Carey VJ. 2013. Software for Computing and Annotating Genomic Ranges. PLOS Comput Biol 9: e1003118.

Lenhard B, Sandelin A, Carninci P. 2012. Metazoan promoters: emerging characteristics and insights into transcriptional regulation. Nat Rev Genet 13: 233–245.

Li X, Noll M. 1994. Compatibility between enhancers and promoters determines the transcriptional specificity of gooseberry and gooseberry neuro in the Drosophila embryo. EMBO J 13: 400–406.

Lizio M, Abugessaisa I, Noguchi S, Kondo A, Hasegawa A, Hon CC, de Hoon M, Severin J, Oki S, Hayashizaki Y, et al. 2019. Update of the FANTOM web resource: expansion to provide additional transcriptome atlases. Nucleic Acids Res 47: D752–D758.

Lizio M, Harshbarger J, Shimoji H, Severin J, Kasukawa T, Sahin S, Abugessaisa I, Fukuda S, Hori F, Ishikawa-Kato S, et al. 2015. Gateways to the FANTOM5 promoter level mammalian expression atlas. Genome Biol 16: 22.

Maricque BB, Chaudhari HG, Cohen BA. 2019. A massively parallel reporter assay dissects the influence of chromatin structure on cis-regulatory activity. Nat Biotechnol 37: 90–95.

McLeay RC, Bailey TL. 2010. Motif Enrichment Analysis: a unified framework and an evaluation on ChIP data. BMC Bioinformatics 11: 165.

Merli C, Bergstrom DE, Cygan JA, Blackman RK. 1996. Promoter specificity mediates the independent regulation of neighboring genes. Genes Dev 10: 1260–1270.

Moudgil A, Wilkinson MN, Chen X, He J, Cammack AJ, Vasek MJ, Lagunas T, Qi Z, Lalli MA, Guo C, et al. 2020. Self-Reporting Transposons Enable Simultaneous Readout of Gene Expression and Transcription Factor Binding in Single Cells. Cell 182: 992–1008.e21.

Ohtsuki S, Levine M, Cai HN. 1998. Different core promoters possess distinct regulatory activities in the Drosophila embryo. Genes Dev 12: 547–556.

Pagès H. 2020. BS genome: Software infrastructure for efficient representation of full genomes and their SNPs. R package version 1580.

Parry TJ, Theisen JWM, Hsu J-Y, Wang Y-L, Corcoran DL, Eustice M, Ohler U, Kadonaga JT. 2010. The TCT motif, a key component of an RNA polymerase II transcription system for the translational machinery. Genes Dev 24: 2013–2018.

Prud’homme B, Gompel N, Carroll SB. 2007. Emerging principles of regulatory evolution. Proc Natl Acad Sci 104: 8605–8612.

Qi Z, Wilkinson MN, Chen X, Sankararaman S, Mayhew D, Mitra RD. 2017. An optimized, broadly applicable piggyBac transposon induction system. Nucleic Acids Res 45: e55.

Roy AL, Singer DS. 2015. Core promoters in transcription: old problem, new insights. Trends Biochem Sci 40: 165–171.

Sharpe J, Nonchev S, Gould A, Whiting J, Krumlauf R. 1998. Selectivity, sharing and competitive interactions in the regulation of Hoxb genes. EMBO J 17: 1788–1798.

Wilkerson MD, Hayes DN. 2010. ConsensusClusterPlus: a class discovery tool with confidence assessments and item tracking. Bioinformatics 26: 1572–1573.

Xiao JY, Hafner A, Boettiger AN. 2021. How subtle changes in 3D structure can create large changes in transcription ed. J. Dekker. eLife 10: e64320.

Yang C, Bolotin E, Jiang T, Sladek FM, Martinez E. 2007. Prevalence of the initiator over the TATA box in human and yeast genes and identification of DNA motifs enriched in human TATA-less core promoters. Gene 389: 52–65.

Yoshida J, Akagi K, Misawa R, Kokubu C, Takeda J, Horie K. 2017. Chromatin states shape insertion profiles of the piggyBac, Tol2 and Sleeping Beauty transposons and murine leukemia virus. Sci Rep 7: 43613.

Zabidi MA, Arnold CD, Schernhuber K, Pagani M, Rath M, Frank O, Stark A. 2015. Enhancer–core-promoter specificity separates developmental and housekeeping gene regulation. Nature 518: 556–559.

